# DDRKOL: A Focused CRISPR Library for Systematic Identification of DNA Damage Response Dependencies in Glioblastoma

**DOI:** 10.64898/2026.09.11.750882

**Authors:** Canan Bayraktar-Odabas, Ipek Kok, Altar Ozbiyik, Alisan Kayabolen, Ali Cenk Aksu, A. Humeyra Dur Karasayar, Ilknur Sur-Erdem, Ibrahim Kulac, Tugba Bagci-Onder

**Author notes:** Corresponding author: Tugba Bagci-Onder, Koç University School of Medicine, Istanbul, Türkiye. Contributed equally (Canan Bayraktar-Odabas, Ipek Kok).

## Abstract

**Background:** DNA damage response (DDR) pathways are central regulators of genome maintenance and major determinants of cancer cell survival. The extensive genomic instability and high replicative stress that characterize glioblastoma render tumor cells highly dependent on DDR pathways to preserve genome integrity and sustain proliferation. This reliance creates potential therapeutic vulnerabilities, making the systematic identification of essential DDR genes a promising strategy for uncovering novel therapeutic targets.

**Methods:** We developed <u>D</u>NA <u>D</u>amage <u>R</u>esponse <u>K</u>nock<u>O</u>ut <u>L</u>ibrary (DDRKOL), a custom CRISPR/Cas9 sgRNA library targeting 819 DDR genes with approximately 10 sgRNAs per gene, together with positive (essential), negative (non-essential) and non-targeting controls. Parallel depletion screens were performed in Cas9-expressing U87-MG and A172 cells cultured for 15 population doublings. Hits were prioritized utilizing TCGA and DepMap databases and validated by viability, clonogenic, apoptosis and GFP competition assays. Clinically relevant patient-derived glioblastoma spheroids and an orthotopic xenograft model was employed to characterize the effects of hit genes.

**Results:** Sequencing confirmed near-complete recovery of the designed sgRNAs from the plasmid pool, with uniform representation across the library and complexity preserved through transduction and selection. Essential-gene controls depleted strongly while non-targeting controls remained neutral, confirming screen performance in both cell lines. The screens identified DDR dependencies in each line and defined a shared core composed of 20 genes belonging to homologous recombination, nucleotide excision repair and ATM/DSB signaling pathways. This shared dependency landscape highlighted four high-confidence candidate genes (*TOP2A, CDK1, XRCC6,* and *RAD21*), which were successfully validated across multiple orthogonal assays. These genes displayed grade-associated expression, and their expressions were positively correlated with proliferation markers in TCGA. Individual knockouts reduced viability, colony formation and competitive fitness, induced apoptosis, and impaired growth of patient-derived glioblastoma spheroids. Both genetic depletion and pharmacological inhibition of TOP2A induced S/G2-M cell cycle arrest. In orthotopic xenografts, TOP2A depletion prevented tumor progression, and led to significantly prolonged survival.

**Conclusion:** DDRKOL represents a robust and versatile focused CRISPR platform for systematic functional interrogation of the DDR associated genes. Using glioblastoma, we demonstrate that the library reliably identifies biologically significant and clinically relevant genetic dependencies through multiple orthogonal validation approaches. As a reusable platform rather than a disease-specific tool, DDRKOL can be broadly applied across diverse biological contexts to discover context-dependent DDR vulnerabilities, therapeutic targets, and mechanisms of treatment resistance.

## BACKGROUND

Glioblastoma (GBM) is the most common and lethal primary malignant brain tumor in adults, characterized by rapid growth, diffuse infiltration into normal brain parenchyma, and near-universal recurrence [1]. Despite maximal safe surgical resection followed by radiotherapy with concurrent and adjuvant temozolomide, median survival remains approximately 12-15 months, with a five-year survival rate below 10% [2]. This dismal outlook has remained essentially unchanged for decades, with attempts to establish more effective therapies yielding little improvement in survival over the last several decades, in marked contrast to progress achieved in many other cancer types [3]. New therapeutic strategies and molecular targets are therefore urgently needed.

A major driver of treatment failure in glioblastoma is the tumor’s capacity to repair therapy-induced DNA lesions. Both radiotherapy and temozolomide act primarily by inducing cytotoxic DNA damage, and DNA damage response (DDR) pathways enable tumor cells to repair this treatment-induced damage, contributing to therapeutic resistance in glioblastoma [4]. Beyond acute damage repair, glioblastoma stem-like subpopulations exhibit heightened expression of DDR factors, which contributes to therapy resistance and disease relapse [3]. DDR signaling is also constitutively engaged because of oncogene-driven replication stress, independent of exogenous genotoxic insult [4]. Classical determinants of chemoresistance, such as MGMT promoter methylation and mismatch repair status account for only part of the clinical variability in treatment response, indicating that unrecognized mechanisms of resistance involving other DDR pathway components remain to be defined [5]. Systematically interrogating the broader DDR network in glioblastoma is therefore expected to reveal both mechanisms of therapy resistance and candidate targets, whose inhibition could re-sensitize tumor cells to standard-of-care treatment [6].

Pooled CRISPR-Cas9 knockout screening has become the method of choice for functional, genome-scale interrogation of gene essentiality and drug sensitivity in cancer, offering higher specificity and lower off-target noise than earlier RNAi-based approaches [7,8]. While genome-wide libraries provide unbiased coverage, their scale imposes substantial cost, sequencing depth, and cell-number requirements, and dilutes statistical power for genes within a specific pathway of interest. Pathway-focused libraries have therefore been developed as a complementary strategy, enabling deeper sgRNA representation per gene and higher-resolution detection of pathway-specific dependencies. We previously generated dedicated CRISPR knockout platforms for other biological compartments, including an epigenetics-focused library whose screening in triple-negative breast cancer and prostate cancer identified novel growth-regulatory epigenetic modifiers [9], and a domain-focused epigenetic library that nominated ASH2L as a context-specific dependency in glioblastoma [10]. However, a comparably comprehensive, DDR-dedicated CRISPR knockout resource, covering the full spectrum of DNA repair and genome-maintenance pathways rather than a handful of candidate genes, has not previously been generated and applied to glioblastoma.

Several individual DDR components have independent precedent as cancer-relevant targets, though their specific contribution to glioblastoma had not been systematically established prior to this work. TOP2A, which resolves topological DNA stress during replication and transcription, is overexpressed in glioma, correlates with proliferative index, and is associated with poor prognosis [11,12]. CDK1, the master regulator of the G2/M transition, is required for mitotic entry, and pharmacological CDK1/2 inhibition arrests glioblastoma cells at the G2-M phase, induces apoptosis and pyroptosis, and reduces tumor volume in xenograft models [13]. XRCC6 (Ku70) initiates non-homologous end joining by binding double-strand break termini, and dysregulation of core NHEJ factors has been repeatedly linked to carcinogenesis, cancer progression, and patient survival across tumor types [14], with NHEJ pathway alterations also documented in glioma [15]. RAD21, a core subunit of the cohesin complex required for sister chromatid cohesion and accurate double-strand break repair, is recurrently overexpressed in solid tumors, where its role in maintaining genomic stability through chromosome segregation, homologous recombination, and telomere preservation appears important in tumor development and has been proposed as a therapeutic target in resistant tumors [16,17]. Despite this cross-cancer precedent, the functional relevance of these genes had not been directly compared or validated within a DDR-wide functional genomics framework.

To address this gap, we designed the DDR Knockout Library, DDRKOL, a custom CRISPR-Cas9 knockout library comprising 8,970 sgRNAs targeting 819 DDR-associated genes spanning all major DNA repair and genome-maintenance pathways, together with positive (essential), negative (non-essential), and non-targeting controls. We applied DDRKOL to parallel screens in two glioblastoma cell lines (U87-MG and A172) to identify genes whose loss selectively impairs GBM cell fitness. Candidate hits were prioritized by cross-referencing screen results with patient survival and expression data and the top-ranked genes were functionally validated across orthogonal assays. This study therefore provides, to our knowledge, the first comprehensive, DDR pathway-wide functional genomic map of genetic dependencies in glioblastoma and nominates TOP2A as a validated therapeutic vulnerability with activity in patient-derived and in vivo GBM models, contributing a resource and a set of prioritized targets for future DDR-directed therapeutic development in this treatment-refractory disease.

## METHODS

### Cell culture

Human glioblastoma cell lines U87-MG, A172, as well as Human Embryonic Kidney 293T cells were purchased from American Type Culture Collection (ATCC). All cell lines were maintained in Dulbecco’s Modified Eagle Medium (DMEM) (Gibco, USA) supplemented with 10% fetal bovine serum (FBS) (Biowest, USA) and 1% Penicillin-Streptomycin (Biowest, USA). Primary glioblastoma cells, GBM8 and GBM4, were kindly gifted by Dr. Hiroaki Wakimoto, Massachusetts General Hospital, Boston, MA [18] and cultured in EF medium consisting of Neurobasal medium (Gibco, USA). Each 500 ml was supplemented with 7.5 mL L-Glutamine (Gibco, USA), 10 mL B27 supplement (Gibco, USA), 2.5 mL N2 supplement (Gibco, USA), 2.5 mL Penicillin-Streptomycin, 500 µL Heparin (Sigma-Aldrich, USA), 100 µL fibroblast growth factor (FGF, 100 µg/mL stock) (PeproTech, USA), and 50 µL epidermal growth factor (EGF, 200 µg/mL stock) (PeproTech, USA). The prepared medium was sterilized by filtration prior to use. All cells were cultured under standard conditions at 37°C in a humidified incubator with 5% CO2. U87-MG and A172 cells were routinely passaged with Trypsin (Gibco, USA). GBM8 cells were collected in suspension, centrifuged at 1200 rpm for 5 minutes, and resuspended in fresh EF medium. Cell viability was routinely assessed using trypan blue exclusion, and seeding densities were adjusted according to the percentage of viable cells.

### Library content of DDRKOL

A total of 819 DDR associated genes were curated based on ENSEMBL annotations. Each gene was targeted by approximately 10 single-guide RNAs (sgRNAs), resulting in 8180 sgRNAs. Sequences were primarily derived from previously established genome-scale libraries, including GeCKO v2 [19] and Brunello [20], and were supplemented with newly designed sgRNAs where necessary. sgRNA design was performed using the Benchling CRISPR design tool, supported by ENSEMBL transcript ID to ensure accurate targeting of coding exons. For each gene, sgRNAs with the highest predicted on-target activity and lowest predicted off-target potential were selected [20]. In addition to DDR-related genes, the library included 35 essential genes (350 sgRNAs) as positive controls, 35 non-essential genes (350 sgRNAs) as negative controls, and 90 non-targeting sgRNAs (approximately 1% of the total library) to serve as internal references. In total, DDRKOL comprised 8970 sgRNAs. Genes and sequences of sgRNAs of DDRKOL are available in **Supplementary Table 1**.

### Cloning of the library

The DDRKOL pooled sgRNA library comprising 8,970 sgRNAs was synthesized as a pooled oligonucleotide library (LC Biosciences, USA). The pooled oligonucleotides were PCR-amplified using Phusion High-Fidelity DNA Polymerase (New England Biolabs, USA), and the resulting amplicons were gel-purified following agarose gel electrophoresis. The pLentiGuide vector (Addgene plasmid #117986) was digested with BsmBI, gel-purified, and assembled with the amplified sgRNA pool using Gibson Assembly Master Mix (New England Biolabs, USA). The assembled library was electroporated into Endura electrocompetent cells (Lucigen, USA) using a MicroPulser (Bio-Rad, USA) according to the manufacturers’ instructions. Following recovery, a small aliquot of the transformed bacteria was serially diluted and plated to estimate library representation, while the remaining culture was expanded overnight in LB medium supplemented with ampicillin. Plasmid DNA was isolated using the NucleoBond Xtra Midi Kit (Macherey-Nagel, Germany). To preserve library complexity, three independent electroporations were performed and pooled after plasmid extraction, resulting in an approximately 1,000× library coverage for subsequent lentiviral production.

### CRISPR/Cas9 screening

Glioblastoma cell lines stably expressing Cas9 (LentiCas9-blast, Addgene #52962) [19] were subjected to three independent biological replicate pooled CRISPR screens. For each biological replicate, 27 × 10⁶ cells were independently transduced with the DDRKOL lentiviral library at a low multiplicity of infection (MOI = 0.3) to ensure single sgRNA integration per cell [19, 20]. Library coverage was maintained at approximately 1000× throughout the screen (27 × 10⁶ cells transduced at MOI = 0.3, yielding ∼9 × 10⁶ puromycin-selected cells per replicate). After 16 h of incubation, the medium was replaced and cells were selected with puromycin (1–2 μg/mL) for 3 days. Following selection, a fraction of cells from each biological replicate was harvested as the baseline sample (Initial Point, IP), while the remaining cells were cultured independently under standard conditions for 30 days (14–16 population doublings). At the endpoint, at least 9 × 10⁶ cells from each replicate were collected (End Point, EP) for genomic DNA extraction (NucleoSpin Tissue Kit, Macherey-Nagel, Germany). sgRNA cassettes were amplified by nested PCR, Illumina adapters were incorporated, and PCR products were gel-purified. Libraries were sequenced on an Illumina platform (Genewiz, USA or GenEra,TR), generating >10 million reads per sample. Sequencing data from the three biological replicates were analyzed using MAGeCK v0.5.9 [21]. The sequences of the primers used for nested PCR and Illumina adapter incorporation are provided in **Supplementary Tables 2–4**.

### Lentiviral packaging and transduction

For viral particle production, 2.5 x 10^6^ HEK293T cells were seeded into 10-cm culture dishes. On the following day, a DNA mixture consisting of 2500 ng transfer vector, 2250 ng Gag-Pol plasmid (psPAX2 for lentivirus or pUMVC for retrovirus), and 225 ng VSV-G was prepared in serum-free DMEM. This DNA mixture was combined with 15 µL PEI reagent diluted in 185 µL serum-free DMEM and incubated at room temperature for 30 min. The transfection mixture was then added dropwise to the cells. After 16h, medium was replaced with 8 mL of fresh growth medium per dish. Viral supernatants were collected at 48 h and 72 h post-transfection, followed by filtration through a 0.45-µm filter [22]. To concentrate viral particles, filtered supernatants were mixed with 5xPEG8000 (Sigma-Aldrich, USA) prepared in PBS at 50% (w/v), yielding a final 1x PEG concentration. The mixture was incubated at 4°C for 1-3 days, then centrifuged at 2500 rpm for 20 min [22]. Resulting viral pellets were resuspended in PBS to obtain a 100x concentrated stock.

For gene ablation using individually cloned sgRNAs, cells were seeded into 6-well plates at a density of 2×10^5^ cells per well. The following day, cells were transduced with lentiviral particles at MOI = 1 in the presence of 10 µg/mL protamine sulfate and selected with puromycin for 3 days.

For lentiviral infection of glioma spheres, cells were resuspended in fresh EF medium at 2×10⁵ – 5×10⁵ cells/mL and incubated with lentiviral particles in the presence of protamine sulfate. Spinfection was performed in 6-well plates at 1000 × g for 60 min at 32°C. Following infection, viral medium was removed, and cells were resuspended in fresh EF medium. Antibiotic selection, when required, was initiated 48h after infection. Forward and reverse oligonucleotide sequences of the individual sgRNAs (two independent guides per gene) used to validate *TOP2A, CDK1, XRCC6,* and *RAD21*, together with the non-targeting control, are provided in **Supplementary Table 5.**

### Cell viability assays

Cell viability was assessed using MTT and CellTiter-Glo (CTG) assays. For MTT assays, cells were seeded at 1000 cells/well in 96-well plates (Sarstedt, Germany) in 100 µL of complete medium. At days 0 and 5, 25 µL of MTT solution (final concentration 0.5 mg/mL) (Cayman, China) was added per well, and plates were incubated for 1-4h at 37°C. Following incubation, the medium was carefully removed and 100 µL DMSO was added to dissolve the formazan. After shaking for 1 min, absorbance was measured at 570 nm using a Synergy H1 plate reader (Biotek, USA). For patient-derived primary GBM8 and GBM4 cells, viability was determined using the CTG Luminescent Cell Viability Assay (Promega, USA). Primary cells were seeded in black-walled 96-well plates at 2000 cells/well in 100 µL medium. At days 0 and 5, CTG reagent was added directly to wells, plates were shaken for 2 min and incubated for 8 min at room temperature. Luminescence, reflecting intracellular ATP content, was measured using a Synergy H1 plate reader. For CRISPR/Cas9 assays, cells were seeded for cell viability at post transduction day (PT) 5.

### Colony formation assay

Cells subjected genetic knockout were seeded at 1500/2000 cells per well in 6-well plates (Sarstedt, Germany) at PT5 and were grown under standard culture conditions for 10-14 days. Following incubation, the medium was removed, cells were washed once with PBS, and fixed with cold methanol at room temperature for 15 min. After an additional PBS wash, colonies were stained with 0.5% crystal violet solution for 45 min. Plates were rinsed gently with water, air-dried and imaged. Quantification of colony number and coverage area was performed using ImageJ software.

### GFP-based competition assay

Cas9-expressing U87-MG and A172 cells were transduced with pBabe-Hygro-GFP (Addgene #61215) and selected with 150 µg/mL Hygromycin. While Cas9-only cells were transduced with NT, Cas9-GFP-expressing cells were transduced with either NT or sgRNA constructs. Following puromycin selection, cells were harvested at PT5. Cas9 and Cas9-GFP cells were combined at a 1:1 ratio, with half of the mixture plated in triplicates and the remaining cells collected for Day 0 baseline measurements. For subsequent analyses, samples were washed with PBS, and GFP signals were measured every 4 days using a CytoFlex flow cytometer (Beckman Coulter, USA).

### Cell Cycle analysis (PI staining)

Cell cycle distribution was evaluated by PI staining and flow cytometry. At PT6 and PT9, cells were collected, washed with PBS, and fixed dropwise in cold 70% ethanol while vortexing. Samples were stored at 4°C for at least 30 min. Fixed cells were centrifuged and resuspended in 100 µL PBS containing RNase A (100 µg/mL, final concentration) and PI (50 µg/mL). Samples were incubated at 37°C for 30 min in the dark. Following gentle vortexing, DNA content was analyzed using a CytoFlex flow cytometer (Beckman Coulter, USA) on the PE channel.

### Apoptosis analysis (Annexin V-FITC/PI staining)

Apoptosis was assessed using Annexin V-FITC/PI staining followed by flow cytometry. At PT6 and PT9, cells were collected and washed with PBS and resuspended in 100 µL of 1X Annexin Binding Buffer. Annexin V-FITC (2.5 µL) and PI (2.5 µL) were added to each sample, gently mixed, and incubated at room temperature for 30 min in the dark. After staining, 400 µL of 1X Annexin Binding Buffer was added to each tube, and samples were analyzed using a CytoFlex flow cytometer (Beckman Coulter, USA) with FITC and PE channels.

### Western Blot

Cell pellets were lysed directly in NP-40 buffer (1% NP-40, 50 mM Tris, 250 mM NaCl, 1x EDTA, 0.02% NaN₃, 1 mM PMSF, and 1x protease inhibitor cocktail) and incubated on ice for 30 min. Lysates were centrifuged at maximum speed for 10 min at 4°C and the supernatants were collected. For nuclear and cytosolic fractionation, cells were washed with PBS and lysed in cytosolic lysis buffer (10 mM HEPES, pH 7.9, 10 mM KCl, 0.1 mM EDTA, 0.4% NP-40, and protease inhibitors) for 15 min on ice [23]. The homogenate was then centrifuged at 3000 × g for 3 min to sediment nuclei, and the supernatant was resedimented at 3000 × g for 5 min; the resulting supernatant was collected as the cytosolic fraction. The nuclear pellet was washed with cytosolic lysis buffer, resuspended in nuclear lysis buffer (20 mM HEPES, pH 7.9, 0.4 M NaCl, 1 mM EDTA, 10% glycerol, and protease inhibitors), sonicated on ice, and centrifuged at 15,000 × g for 5 min; the resulting supernatant was collected as the nuclear fraction. For protein quantification, the Pierce BCA Protein Assay Kit (Thermo Scientific, USA) was used. Equal amounts of protein (20–30 µg) were denatured in 4x Laemmli buffer (Bio-Rad, USA) supplemented with 1:10 β-mercaptoethanol by boiling at 95°C for 10 min. Samples were resolved on precast 4–12% Mini-PROTEAN TGX gels (Bio-Rad, USA) and subsequently transferred onto PVDF membranes using the Trans-Blot Turbo Transfer System (Bio-Rad, USA). Membranes were blocked for 1h at room temperature with either 5% non-fat dry milk or 5% bovine serum albumin (BSA) prepared in TBS-T. Blots were then incubated with primary antibodies overnight at 4°C, followed by three washes with TBS-T. HRP-conjugated secondary antibodies were applied for 1h at room temperature, and signals were visualized using the Odyssey FC Imaging System (LI-COR Biosciences, USA). The list of primary and secondary antibodies used in these experiments is provided in **Supplementary Table 6**.

### qRT-PCR

RNA was extracted using commercially available kit (Macherey-Nagel, Germany) and a total of 500-1000 ng RNA was reverse transcribed into cDNA using the M-MLV Reverse Transcriptase kit (Invitrogen, USA). Quantitative real-time PCR (qRT-PCR) was carried out on a LightCycler 480 Instrument II (Roche, Switzerland) with each reaction performed in triplicate. The cycling program was set as follows: an initial denaturation at 95°C for 5 min, followed by 45 cycles of 95°C for 10s, 60°C for 30s, and 72°C for 30s. GAPDH was used as the internal reference gene, and relative expression levels were calculated using the 2^ΔΔCT^ method. Primer sequences used for qRT-PCR are provided in **Supplementary Table 7**.

### Sphere Size and Volume Analysis

Sphere size was quantified from phase-contrast microscopy images using Fiji/ImageJ software (USA). Before measurement, the image scale was calibrated using the corresponding microscope scale bar. For each sphere, the diameter was measured by drawing a straight line across the widest part of the sphere using the Straight Line tool. Sphere volume was then estimated from the measured diameter by assuming an approximately spherical geometry. Measurements were performed using images acquired 7 days after the end of antibiotic selection for GBM4 cultures and 18 days after the end of antibiotic selection for GBM8 cultures.

### Drug treatment and IC50 determination

Etoposide (Cayman Chemical, #12092) and topotecan (Prestwick Chemical Library, France) were dissolved in DMSO to prepare 10 mM stock solutions, which were stored at −80°C and protected from light. For dose–response experiments, U87-MG cells were seeded at 1,000 cells/well in 96-well plates (Sarstedt, Germany) in 100 µL complete medium and allowed to adhere overnight. Cells were then treated with a serial dilution series of etoposide (0.1–40 µM) or topotecan (0.016–2 µM), together with a vehicle control (DMSO, matched to the highest drug concentration, ≤0.1% v/v) and an untreated control. After 48h of exposure, viability was assessed by MTT assay as described above, and absorbance was measured at 570 nm using a Synergy H1 plate reader (Biotek, USA). Viability values were normalized to DMSO-treated controls, and IC50 values were calculated in GraphPad Prism 10.0 (USA) by fitting log-transformed drug concentrations to a four-parameter variable-slope nonlinear regression model. For cell cycle experiments, U87-MG cells were seeded in 6-well plates at 100.000 cells/well and treated the following day with etoposide or topotecan at concentrations corresponding to their respective IC50 values determined. Vehicle controls (DMSO) and untreated controls (0 µM) were included for each drug. After 48h of treatment, cells were harvested and processed for propidium iodide staining and flow cytometry as described in the cell cycle analysis section. For comparison with genetic depletion, cells transduced with TOP2A-targeting or non-targeting sgRNAs were analyzed in parallel at PT6 and PT9 without drug treatment.

### In vivo tumor growth assay

U87-MG cells stably expressing Firefly luciferase (FLuc) and mCherry were generated by lentiviral transduction as described [10]. For functional validation, sgRNA was cloned into pLCV2 vector and introduced into cells via lentiviral transduction. Five days post-transduction, 1.5×10^5^ cells in 7 µL PBS were injected intracranially into the left frontal lobe of 6-8-week-old NOD/SCID mice using a stereotactic frame (Coordinates: 2 mm lateral and 2 mm caudal to bregma, 2 mm depth from dura). Short-term anesthesia was achieved with intraperitoneal ketamine (80-100 mg/kg) and xylazine (5-10 mg/kg). Tumor growth was monitored weekly for up to four weeks using the IVIS Lumina III Imaging system (Perkin-Elmer, USA). Bioluminescence imaging was performed following intraperitoneal administration of D-luciferin (150 µg/g body weight). During imaging, mice were anesthetized with isoflurane. Tumor volume was quantified from FLuc signal intensity, while animal weight and behavior were monitored throughout the study.

Brains were harvested, sectioned coronally from anterior to posterior, and processed on an automated tissue processor (Tissue-Tek CIP 6 AI, Sakura, Japan). Following dehydration in graded ethanol (70– 100%), xylene clearing, and paraffin embedding, 5 µm sections were cut by microtome. Sections were deparaffinized, rehydrated through graded alcohols, and stained with hematoxylin and eosin according to the manufacturer’s protocol (Leica, Germany). All sectioning and staining were performed by the Department of Pathology, Koç University Hospital.

### Clinical dataset analyses

Clinical characterization of candidate genes was performed using publicly available glioma datasets from The Cancer Genome Atlas (TCGA) GBM and GBM-LGG (GBM-low grade glioma) cohorts [24]. Kaplan-Meier survival analysis and mRNA expression comparisons across histological subtypes (GBM, astrocytoma, oligodendroglioma, and oligoastrocytoma) were performed using the GlioVis data portal [25], which provided hazard ratios with 95% confidence intervals, log-rank p-values, and Wilcoxon p-values for survival curves, and pairwise t-tests with Bonferroni correction for cross-subtype expression comparisons. Differential mRNA expression between GBM tumor samples and normal brain cortex, together with Pearson correlation analyses between candidate gene expression and MKI67 (Ki-67) expression across GBM samples, were performed using GEPIA2 [26], which integrates RNA-sequencing data from TCGA tumor samples and matched normal tissue from the Genotype-Tissue Expression (GTEx) project. Log2 fold changes and adjusted p-values for tumor-versus-normal comparisons, and Pearson correlation coefficients with associated significance levels for gene-gene correlation analyses, were derived directly from these platforms’ outputs.

### Statistical analysis

Statistical analyses were performed using GraphPad Prism 10. Data are presented as mean ± SEM unless otherwise indicated. The statistical tests used for each experiment are specified in the corresponding figure legends.

## RESULTS

### Design and quality assessment of the DDRKOL CRISPR knockout library

To enable focused, high-resolution functional interrogation of the DDR in glioblastoma, we custom designed a CRISPR-Cas9 knockout library (DDRKOL) targeting the DDR gene network at high sgRNA density (**Fig. 1A**). The library comprised 8,970 sgRNAs directed against 819 DDR-associated genes, together with positive (essential), negative (non-essential), and non-targeting controls, with each gene targeted by 10 independent sgRNAs to maximize statistical power for per-gene depletion cells (**Fig. 1B**) [7,19, 20]. Curated DDR genes spanned all major DNA repair and genome-maintenance pathways, with the largest representation in homologous recombination (21.5%), ATM/DSB signaling (14.1%), and nucleotide excision repair (8.6%), reflecting the breadth of the DDR network captured by the library (**Fig. 1B**). At the pathway level, coverage was high across nearly all functional categories: core pathways including editing and processing nucleases, non-homologous end joining, and Fanconi anemia were targeted at 100% gene-level coverage, and homologous recombination, DNA polymerases, base excision repair, and nucleotide excision repair each exceeded 90% (**Fig. 1D**).

**Figure 1.**
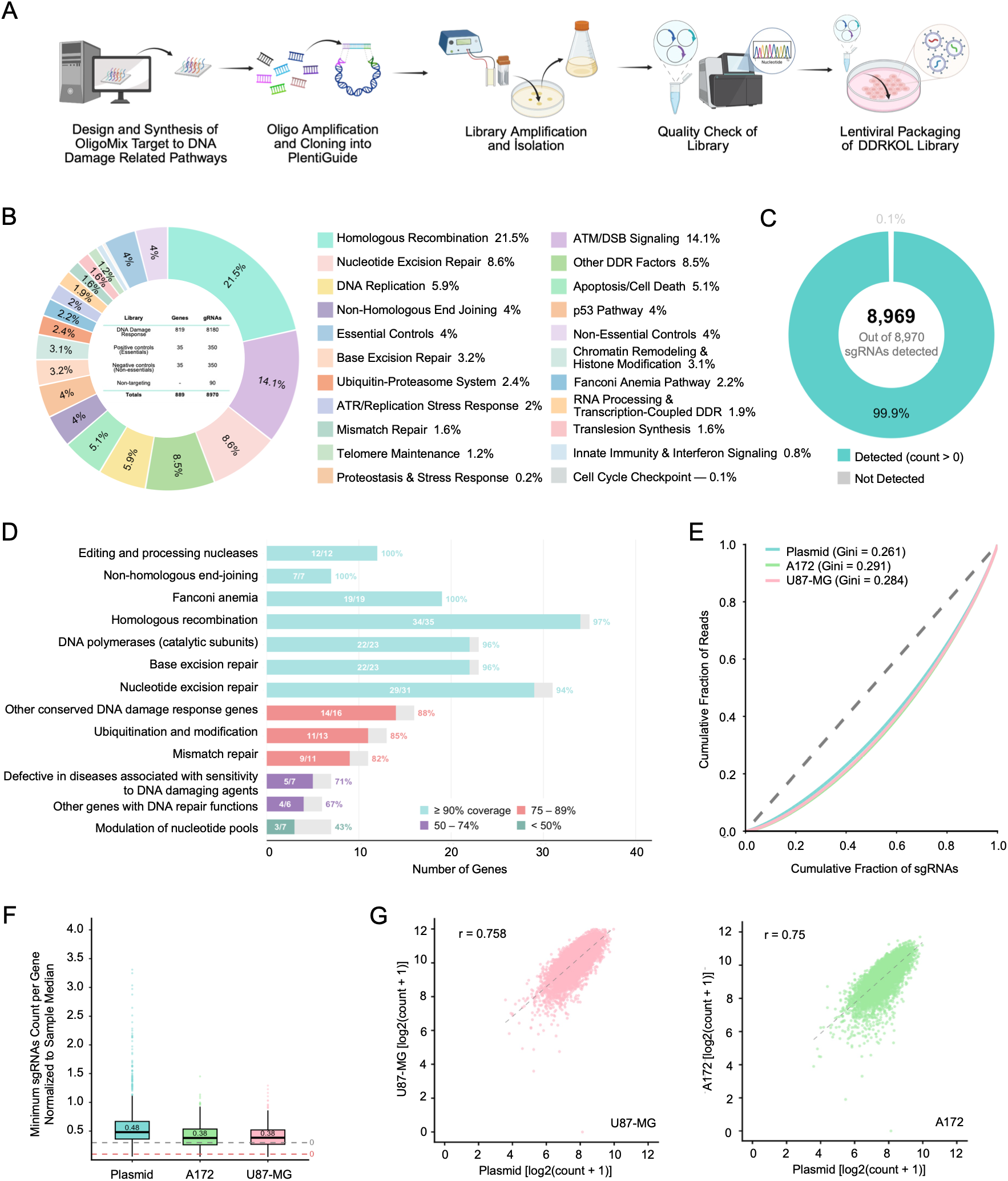
Design, composition, and quality assessment of the DDRKOL CRISPR knockout library. **A)** Schematic overview of the DDRKOL library generation workflow. Oligonucleotide pools targeting DNA damage-related pathways were designed and synthesized via Oligomix, amplified, and cloned into the plentiGuide backbone. The resulting library was amplified, isolated, and subjected to quality control prior to lentiviral packaging. **B)** Functional composition of the custom DNA damage response knockout library (DDRKOL). The library consists of 8,970 sgRNAs targeting 819 DDR-associated genes together with positive (essential), negative (non-essential), and non-targeting controls. DDR genes were categorized into major DNA repair and genome maintenance pathways according to their primary biological functions. **C)** Donut chart showing plasmid library sequence representation. Of the 8,970 designed sgRNAs, 8,969 (99.9%) were detected with at least one read following next-generation sequencing of the plasmid pool. **D)** Horizontal bar chart depicting the DDR pathway coverage of the DDRKOL library. Bars represent the number of genes targeted per pathway, color-coded by coverage level: >95% (teal), 75–95% (salmon), 50–74% (blue), and <50% (purple). Gray bars indicate the total number of genes annotated per pathway. **E)** Lorenz curves depicting cumulative sgRNA read distribution for the plasmid library, A172, and U87-MG cell lines. Gini coefficients (Plasmid = 0.261, A172 = 0.291, U87-MG = 0.284) indicate highly uniform sgRNA representation comparable to published genome-wide libraries. The dashed diagonal represents perfect equality. **F)** Boxplot of per-gene relative minimum sgRNA count, defined as the minimum sgRNA count among the 10 sgRNAs targeting each gene normalized to the sample median. Median values are indicated above each box. Red and gray dashed lines denote relative minimum thresholds of 0.1 and 0.3, respectively, indicating adequate and good per-gene representation. **G)** Scatter plots of log₂-transformed normalized sgRNA counts comparing the plasmid library to U87-MG (left, pink) and A172 (right, green) cell lines. Pearson correlation coefficients (r = 0.758 and r = 0.750, respectively) confirm strong concordance between the plasmid reference and transduced cell populations. Dashed lines indicate linear regression fits.

We next assessed the technical quality of the synthesized library. Next-generation sequencing of the plasmid pool detected 8,969 of the 8,970 designed sgRNAs (99.9%) with at least one read, confirming near-complete representation with negligible dropout during oligonucleotide synthesis and cloning (**Fig. 1C**). To evaluate the uniformity of sgRNAs representation, we computed the Lorenz curve and Gini coefficient for the plasmid library and for each transduced glioblastoma cell line [27]. Read distributions were tightly balanced, with Gini coefficients of 0.261 for the plasmid library, 0.284 for U87-MG, and 0.291 for A172 (**Fig. 1E**), and frequency histograms of log2-normalized counts were unimodal and symmetric across all three samples, consistent with successful lentiviral transduction and even library propagation (**Supp. Fig.1**). Critically, per-gene representation was preserved even at the most stringent measure: the relative minimum sgRNA count, defined as the lowest-abundance sgRNA among the ten targeting each gene, normalized to the sample median, had a median of 0.48 in the plasmid library and 0.38 in both cell lines, exceeding the 0.1 and 0.3 thresholds indicative of good per-gene coverage and confirming that few genes were left under-represented by chance dropout of individual guides (**Fig. 1F**).

Finally, to confirm that library complexity was faithfully maintained through transduction and selection, we compared log2-normalized sgRNA counts between the plasmid pool and each cell line. Both glioblastoma lines showed strong concordance with the input library (Pearson r = 0.758 for U87-MG and r = 0.750 for A172; **Fig. 1G**), indicating that the abundance structure of the library was preserved during infection and puromycin selection rather than distorted by bottlenecking. Together, these data establish DDRKOL as a well-constructed, deeply representative, DDR-focused screening resource, and confirm that both U87-MG and A172 support high-quality screening with this library.

### A focused CRISPR screen identifies DDR dependencies in glioblastoma

To identify DDR genes required for glioblastoma fitness, we performed parallel negative-selection CRISPR screens in two Cas9-expressing cell lines, U87-MG and A172, with high Cas9 activity confirmed by a GFP reporter assay (**Fig. 2A; Supp. Fig. 2A**). Cells were transduced with the DDRKOL library at low MOI, selected with puromycin, and sampled at InitialPoint and after 15 population doublings (PDL) (EndPoint) for sgRNA sequencing (**Fig. 2A**). Both screens behaved as expected for a high-quality dropout experiment: positive-control essential sgRNAs were strongly depleted at EndPoint, whereas both non-essential and non-targeting controls remained centered near neutrality (**Fig. 2B**) [7,28]. Gene-level depletion was quantified with MAGeCK [21], which ranked known essential genes at the extreme depleted tail (**Supp. Fig. 2B**), and significant hits were defined from the volcano-plot distributions (**Fig. 2C**). The most strongly depleted genes were canonical DDR factors, including *PCNA, POLE2, XRCC6, RAD21,* and *TOP2A* in A172, and *PIDD1, TFDP1, UHRF1, RAD21,* and *POLD1* in U87-MG (**Supp. Fig. 2C**).

**Figure 2.**
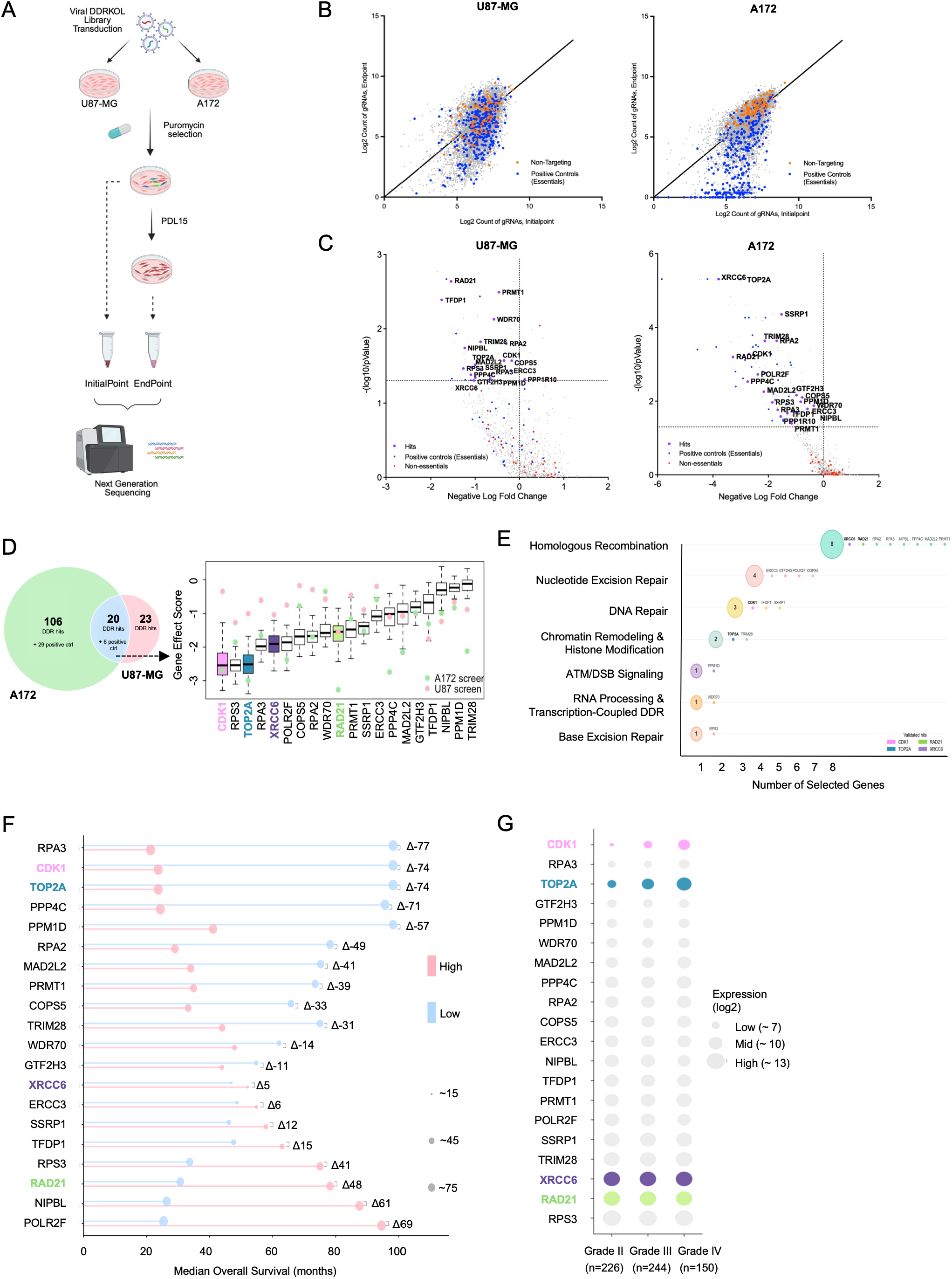
Parallel DDRKOL screens in U87-MG and A172 cells define a shared core of DDR dependencies in glioblastoma. **A)** Schematic of the DDRKOL screen workflow. U87-MG and A172 cells were transduced with the DDRKOL library, selected with puromycin, and harvested at PDL15 (EndPoint) and day 0 (InitialPoint) for next-generation sequencing. **B)** Scatter plots of log₂-normalized sgRNA counts at InitialPoint vs. EndPoint for U87-MG (left) and A172 (right). Positive controls (essentials, blue) and non-targeting controls (orange) are indicated. **C)** Volcano plots of CRISPR screen results for U87-MG (left) and A172 (right). x-axis: negative log₂ fold change; y-axis: −log₁₀(p-value). Screen hits (purple), essential positive controls (blue), and non-essential controls (red) are shown. Labeled genes indicate common hits. Dashed lines indicate significance thresholds. **D)** Identification of common dependencies across glioblastoma cell lines. Left, Venn diagram showing the overlap of significantly depleted DDR genes identified in the A172 and U87-MG CRISPR knockout screens. A total of 20 common DDR hits and 6 shared positive control genes were identified between the two screens. Right, gene effect scores of the shared DDR hits ranked according to their dependency across both cell lines. Colored dots represent individual gene effect scores from the A172 (green) and U87-MG (pink) screens, while boxplots summarize the distribution of gene dependency scores across the DepMap CRISPR dataset. **E)** Bubble plot showing pathway distribution of screen hits. Bubble size reflects number of selected genes per pathway; individual gene dots are shown to the right. **F)** Lollipop plot depicting median overall survival (months) for patients stratified by high vs. low grade glioma of screen hit genes (TCGA). Colored dots indicate direction of effect; values shown to the right. **G)** Dot plot showing expression levels of screen common hit genes across glioma grades II, III, and IV (TCGA). Dot size reflects expression level.

A172 yielded more DDR hits (106) than U87-MG (23), with twenty genes common to both lines representing a high-confidence core of glioblastoma dependencies (**Fig. 2D**). Comparison with genome-wide dependency data from the Cancer Dependency Map (DepMap) [29] showed these shared hits spanned a spectrum from pan-essential genes (e.g., *CDK1, RPS3, TOP2A*) to more selective dependencies (e.g., *XRCC6, RAD21)* (**Fig. 2D**). At the pathway level, dependencies were concentrated in homologous recombination, nucleotide excision repair, and ATM/DSB signaling in both lines, with A172 additionally enriched for DNA replication and apoptosis factors (**Fig. 2E; Supp. Fig. 3A, B**). To capture the breadth of the identified DDR vulnerabilities, four candidate genes (*TOP2A, CDK1, XRCC6,* and *RAD21*) representing distinct functional branches of the DDR network were prioritized for further validation (**Fig. 2E**). These four candidates were selected because they were reproducibly depleted in both cell lines, were supported by multiple independent sgRNAs, and collectively spanned both pan-essential and comparatively selective dependencies across distinct DDR functions. Their prioritization was further supported by patient-level expression and survival associations, enabling subsequent validation to integrate screening robustness, mechanistic diversity, and clinical relevance rather than relying on depletion magnitude alone.

**Figure 3.**
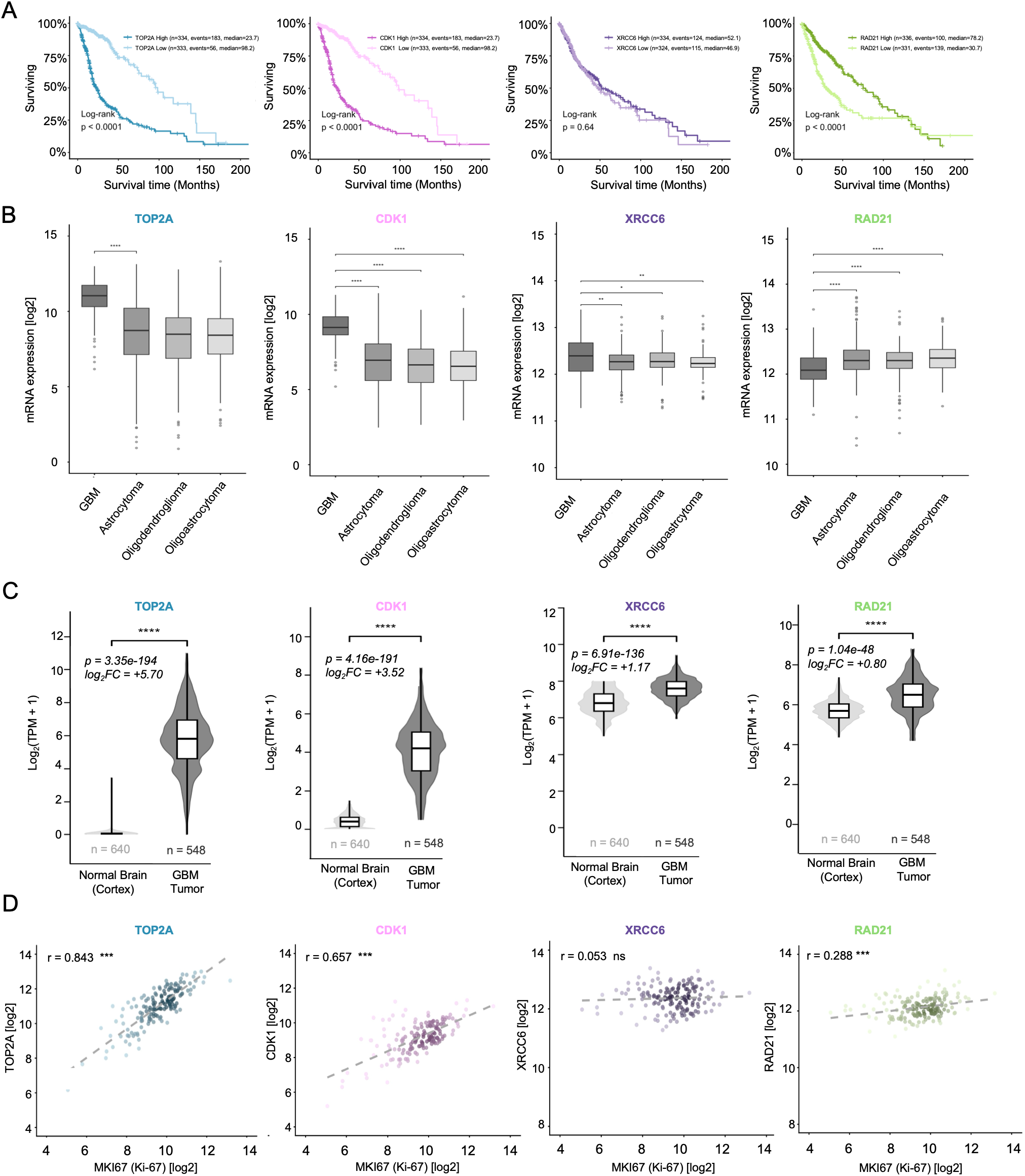
Clinical and transcriptomic characterization of DDRKOL screen candidate genes in glioblastoma. **A)** Kaplan-Meier survival curves for patients stratified by high (red) or low (teal) expression of TOP2A, CDK1, XRCC6, and RAD21 (TCGA GBMLGG dataset). Hazard ratios (HR) with 95% confidence intervals, log-rank p-values, and Wilcoxon p-values are indicated. Number of patients and median survival times are shown in the legend of each plot. **B)** mRNA expression of the four candidate genes across glioma histological subtypes; GBM (n=152), astrocytoma (n=194), oligodendroglioma (n=191), and oligoastrocytoma (n=130) from the TCGA GBMLGG dataset. Statistical significance was assessed by pairwise t-tests with Bonferroni correction; *p < 0.05, **p < 0.01, ***p < 0.001, ns, not significant. **C)** Violin plots depicting log₂-transformed mRNA expression of TOP2A, CDK1, XRCC6, and RAD21 in GBM tumor or normal brain samples. Log₂ fold change and adjusted p-values are indicated above each comparison. **D)** Scatter plots showing Pearson correlation between MKI67 (Ki-67) expression and each candidate gene across samples (TCGA GBMLGG, GBM histology, n=152). Pearson r values and significance levels are indicated; ***p < 0.001, ns, not significant. TOP2A (r = 0.843) and CDK1 (r = 0.657) show strong positive correlation with the proliferation index, while RAD21 (r = 0.288) shows moderate and XRCC6 (r = 0.053) shows no significant correlation.

Finally, integrating the shared hits with The Cancer Genome Atlas (TCGA) patient data [24] showed that most, including *CDK1*, *TOP2A*, and *RAD21*, increased in expression with advancing tumor grade (**Fig. 2F; Supp. Fig. 3C**), and that expression of many was associated with differences in median overall survival (**Fig. 2G**). Together, these screens define a reproducible set of DDR dependencies in glioblastoma, distinguish core from context-specific hits, and nominate *TOP2A, CDK1, XRCC6,* and *RAD21* as clinically correlated candidates for validation.

### Candidate DDR dependencies are clinically linked to glioblastoma

To evaluate the clinical relevance of the four prioritized candidates, we examined their expression and prognostic associations in glioma patient cohorts from TCGA [24]. Kaplan-Meier analysis of GBM and low-grade glioma (LGG) patients stratified by candidate expression revealed strong prognostic associations for three of the four genes: high TOP2A and high CDK1 expression were each associated with markedly shorter overall survival, and RAD21 expression was likewise significantly associated with survival, whereas XRCC6 showed no significant survival association (**Fig. 3A**). This divergence indicated that, although all four genes are functional dependencies in the screen, their expression carries distinct prognostic weight in patients.

We next assessed expression across glioma histological subtypes using the GlioVis data portal [25]. TOP2A and CDK1 were most highly expressed in GBM relative to lower-grade astrocytoma, oligodendroglioma, and oligoastrocytoma, while XRCC6 and RAD21 showed more modest subtype-dependent differences (**Fig. 3B**). Comparison of GBM tumors against normal brain cortex using GEPIA2 [26] confirmed that all four candidates were significantly overexpressed in tumor tissue, though with very different magnitudes: TOP2A and CDK1 showed pronounced upregulation, whereas XRCC6 and RAD21 were more modestly elevated (**Fig. 3C**).

To determine whether these expression patterns reflected tumor proliferation, we correlated each candidate with MKI67 (Ki-67), a canonical marker of proliferating cells [30], across GBM samples (**Fig. 3D**). TOP2A and CDK1 correlated strongly with the proliferation index, and RAD21 showed a weaker but significant correlation, whereas XRCC6 showed no significant correlation. These results indicate that the candidates occupy distinct functional niches in GBM: TOP2A and CDK1 behave as proliferation-coupled, prognostically adverse genes, RAD21 as an intermediate proliferation-associated dependency, and XRCC6 as a proliferation-independent dependency whose essentiality in the screen is not explained by expression level or proliferative state. Together, these analyses define TOP2A and CDK1 as the most clinically compelling candidates while highlighting mechanistically distinct roles for RAD21 and XRCC6.

### Individual knockout of TOP2A, CDK1, XRCC6, and RAD21 impairs glioblastoma cell fitness

To validate the four prioritized candidates, we first examined their sgRNA-level behavior in the primary screen. Slope graphs of individual sgRNA abundance from InitialPoint to EndPoint showed consistent depletion of the majority of guides targeting TOP2A, CDK1, XRCC6, and RAD21 in both U87-MG and A172 (**Fig. 4A**), confirming that the gene-level dropout signal was driven by multiple independent sgRNAs rather than a single outlier guide. We then individually knocked out each candidate using two independent sgRNAs (g1, g2) in both cell lines (**Fig. 4B**), with knockout efficiency confirmed at the protein level by Western Blot (**Fig. 4E**) and at the transcript level by qRT-PCR (**Supp. Fig. 4**). Non-targeting sgRNA (NT) as a negative control and the ribosomal gene RPL9 as a positive (lethal) control throughout.

**Figure 4.**
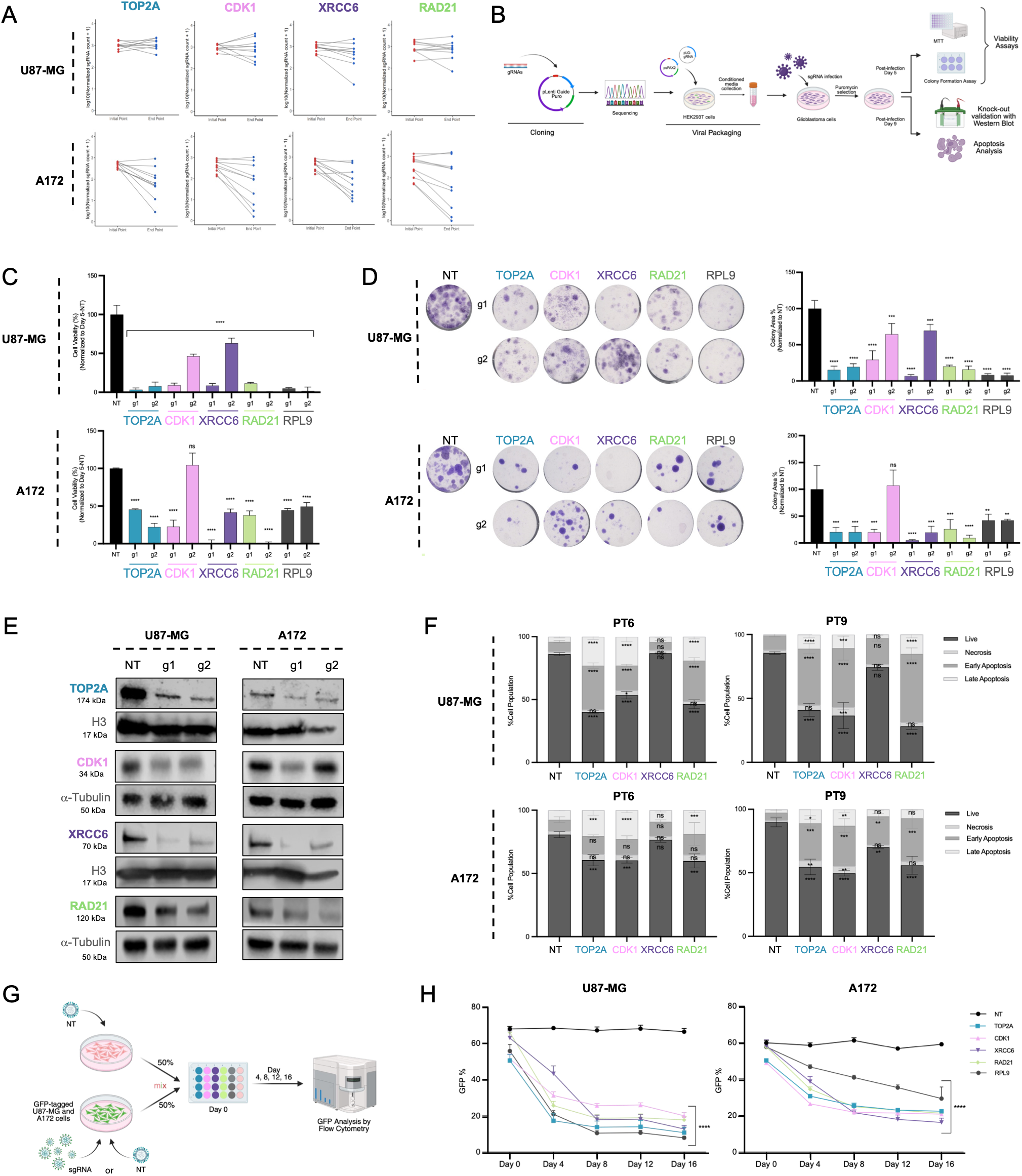
Individual knockout of TOP2A, CDK1, XRCC6, and RAD21 impairs glioblastoma cell fitness. **A)** Slope graphs depicting log₂-normalized sgRNA counts at InitialPoint and EndPoint for TOP2A, CDK1, XRCC6, and RAD21 in U87-MG (top) and A172 (bottom). Each line represents one sgRNA; red = depleted, blue = enriched. **B)** Schematic of individual gene validation workflow. **C)** MTT viability assay in U87-MG cells following knockout of TOP2A, CDK1, XRCC6, RAD21 and RPL9 (n=3, mean ± SEM). **D)** Representative images (left) and quantification (right) of clonogenic survival assays in U87-MG (top) and A172 (bottom) cells. Colony area normalized to NT control (mean ± SEM). For panels C and D, statistical significance was determined using one-way ANOVA followed by Dunnett’s multiple comparisons test, with NT serving as the control group. **E)** Western blots confirming knockout efficiency for TOP2A, CDK1, XRCC6 and RAD21 in U87-MG and A172 cells. NT = non-targeting control; g1, g2 = independent sgRNAs. **F)** Stacked bar charts showing cell fate distribution (live, necrosis, early apoptosis, late apoptosis) by Annexin V/PI staining at PT6 and PT9 in U87-MG (top) and A172 (bottom). Statistical analyses for panel F were performed using two-way ANOVA followed by Šídák’s multiple comparisons test to compare knockout and control groups within each cell fate and time point. **G)** Schematic workflow of GFP-based Competition Assay. **H)** GFP competition assay tracking the proportion of GFP⁺ cells over time in U87-MG (top) and A172 (bottom) following knockout of indicated genes. Data were summarized by calculating the area under the curve (AUC) for each biological replicate. AUC values were analyzed using one-way ANOVA followed by Dunnett’s multiple comparisons test, with NT as the control group. Data are presented as mean ± SEM. Statistical significance is indicated as follows: ns, not significant; *p* < 0.05 (\**); p < 0.01 (**); p < 0.001 (\*\*\**); *p < 0.0001 (****)*.

Knockout of each candidate markedly reduced cell viability. In MTT assays, depletion of TOP2A, CDK1, XRCC6, and RAD21 significantly decreased viability relative to NT in both cell lines, in most cases approaching the effect of the essential-gene control RPL9 (**Fig. 4C**). Long-term clonogenic survival, a stringent measure of cell fitness [31], was even more severely impaired: colony formation was significantly suppressed by knockout of all four genes in both U87-MG and A172, with most sgRNAs reducing colony area to a small fraction of the NT control (**Fig. 4D**). The concordance between the acute (MTT) and long-term (clonogenic) readouts, across two independent sgRNAs per gene, established that the fitness defects were robust and on target.

To determine how loss of each gene compromises glioblastoma cell fitness, we assessed cell death by Annexin V/PI staining at two post-transduction time points (PT6 and 9). Knockout of each candidate progressively shifted the cell population from viable toward early and late apoptotic fractions relative to NT, with the effect increasing from PT6 to PT9 in both cell lines (**Fig. 4F**), indicating that candidate loss triggers apoptotic cell death rather than a purely cytostatic arrest. Finally, we quantified competitive fitness directly using a GFP-based competition assay, in which knockout cells are co-cultured with control cells and their relative proportion tracked over time (**Fig. 4G**). Cells depleted of TOP2A, CDK1, XRCC6, and RAD21 were progressively outcompeted, with the GFP+ fraction declining steadily over 16 days in both U87-MG and A172, closely mirroring the depletion kinetics of the RPL9 essential control (**Fig. 4H**). Together, these orthogonal assays consistently confirm that each of the four screen candidates is individually required for glioblastoma cell fitness, validating the primary screen and establishing TOP2A, CDK1, XRCC6, and RAD21 as bona fide glioblastoma dependencies.

### Candidate knockout impairs growth of patient-derived glioblastoma spheroids

Because established serum-cultured cell lines can diverge genetically and phenotypically from the tumors they originate from, patient-derived glioblastoma spheroid cultures, which more faithfully recapitulate the genotype, expression profile, and biology of primary tumors, provide a more clinically representative validation setting [32]. We therefore tested whether loss of the four candidates impaired growth in GBM8, a patient-derived glioblastoma model grown as three-dimensional spheroids. CRISPR mediated knockout of TOP2A, CDK1, XRCC6, and RAD21 each significantly reduced spheroid size relative to the NT control, as did the essential-gene control RPL9 (**Fig. 5A**). This effect was evident morphologically: NT spheroids formed large, compact, well-defined structures, but knockout of the candidate genes yielded small, poorly organized, or dispersed cell aggregates that largely failed to form intact spheroids (**Fig. 5B**). To complement the morphological analysis with a metabolic readout, we performed a luminescence-based viability assay at PT9. Knockout of TOP2A, CDK1, and RAD21 significantly reduced viability relative to NT, consistent with the spheroid-volume data, whereas XRCC6 knockout produced a more modest, non-significant reduction in this readout (**Fig. 5C**). To confirm that these effects were not restricted to a single patient-derived model, we repeated the viability assay in a second independent line, GBM4. Here, knockout of all four candidates, including XRCC6, significantly reduced viability relative to NT (**Supp. Fig. 5**), reinforcing the generality of the dependency across patient-derived backgrounds. The comparatively weaker and more variable effect of XRCC6 knockout in the GBM8 viability assay, despite its clear reduction of spheroid size, is consistent with the distinct, proliferation-independent behavior of XRCC6 observed in the clinical and transcriptomic analyses (**Fig. 3**). Overall, these results extend the validation of TOP2A, CDK1, XRCC6, and RAD21 from established cell lines to clinically relevant patient-derived glioblastoma models, strengthening their credibility as therapeutic targets.

**Figure 5.**
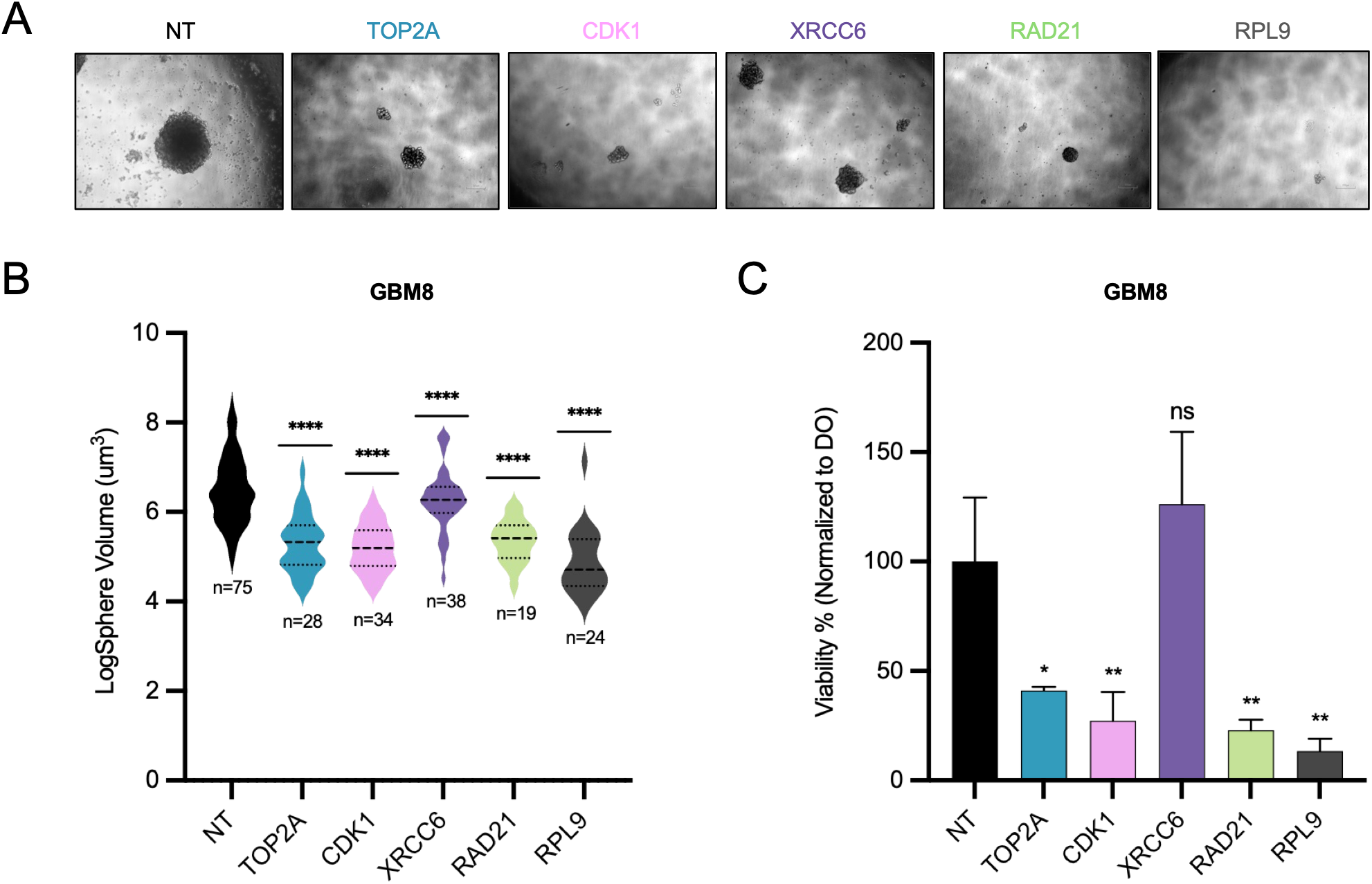
Candidate gene knockout impairs growth of patient-derived glioblastoma spheroids. **A)** Representative bright-field images of GBM8 spheroids following knockout of the indicated genes. Spheroid morphology and growth were assessed under indicated sgRNA conditions to evaluate the effects of candidate DDR gene depletion on tumorsphere formation and maintenance. Scale bars, 200 μm**. B)** Quantification of spheroid size following CRISPR-mediated knockout of the indicated genes in GBM8 spheroids. Spheroid volumes were measured and log-transformed for analysis. The number of spheroids analyzed for each condition is indicated. **C)** Cell Titer Glo viability assay for PT9 in GBM8 cells following indicated knockouts. Log-transformed spheroid volume data and CellTiter-Glo (CTG) assay data were analyzed using one-way ANOVA followed by Dunnett’s multiple comparisons test, with NT as the control group. Data are presented as mean ± SEM. Statistical significance was defined as ns, not significant; *p* < 0.05 (\**), p < 0.01 (**), p < 0.001 (\*\*\**), and *p < 0.0001 (****)*.

### TOP2A depletion arrests cell cycle progression and abrogates orthotopic glioblastoma growth

To place TOP2A in a broader functional context, we performed co-dependency analysis using genome-wide CRISPR data from the Cancer Dependency Map [29]. The genes most strongly correlated with TOP2A dependency were dominated by cell cycle and DNA repair factors, led by NCAPH2, ESPL1, TOP3A, RMI1, KIF23, and RACGAP1, and including multiple condensin complex subunits (**Supp. Fig. 6A**). Consistent with a role in the most aggressive disease context, TOP2A expression in the TCGA GBMLGG cohort was significantly higher in IDH-wildtype GBM (n=55) than in IDH-mutant non-codeleted (n=141) or IDH-mutant codeleted (n=85) tumors (**Supp. Fig. 6B**).

Because TOP2A is the target of clinically used topoisomerase inhibitors [33], we asked whether pharmacological inhibition phenocopies genetic depletion. U87-MG glioblastoma cells were sensitive to both the TOP2 inhibitor etoposide (IC50 = 1.580 µM) and the TOP1 inhibitor topotecan (IC50 = 0.322 µM) (**Fig. 6A**). Cell cycle analysis revealed that TOP2A knockout progressively depleted the G0/G1 fraction relative to non-targeting controls, from approximately 73% to 61% at PT6 and from 87% to 66% at PT9, with a corresponding accumulation in S and G2/M (**Fig. 6B**). Drug treatment produced the same directional shift far more acutely: etoposide collapsed the G0/G1 population to under 10% with most cells accumulating in S phase, while topotecan produced a more moderate reduction relative to DMSO controls (**Fig. 6B**). Genetic and pharmacological loss of TOP2A function therefore converge on the same phenotype, failure to complete S phase and progress through mitosis.

**Figure 6.**
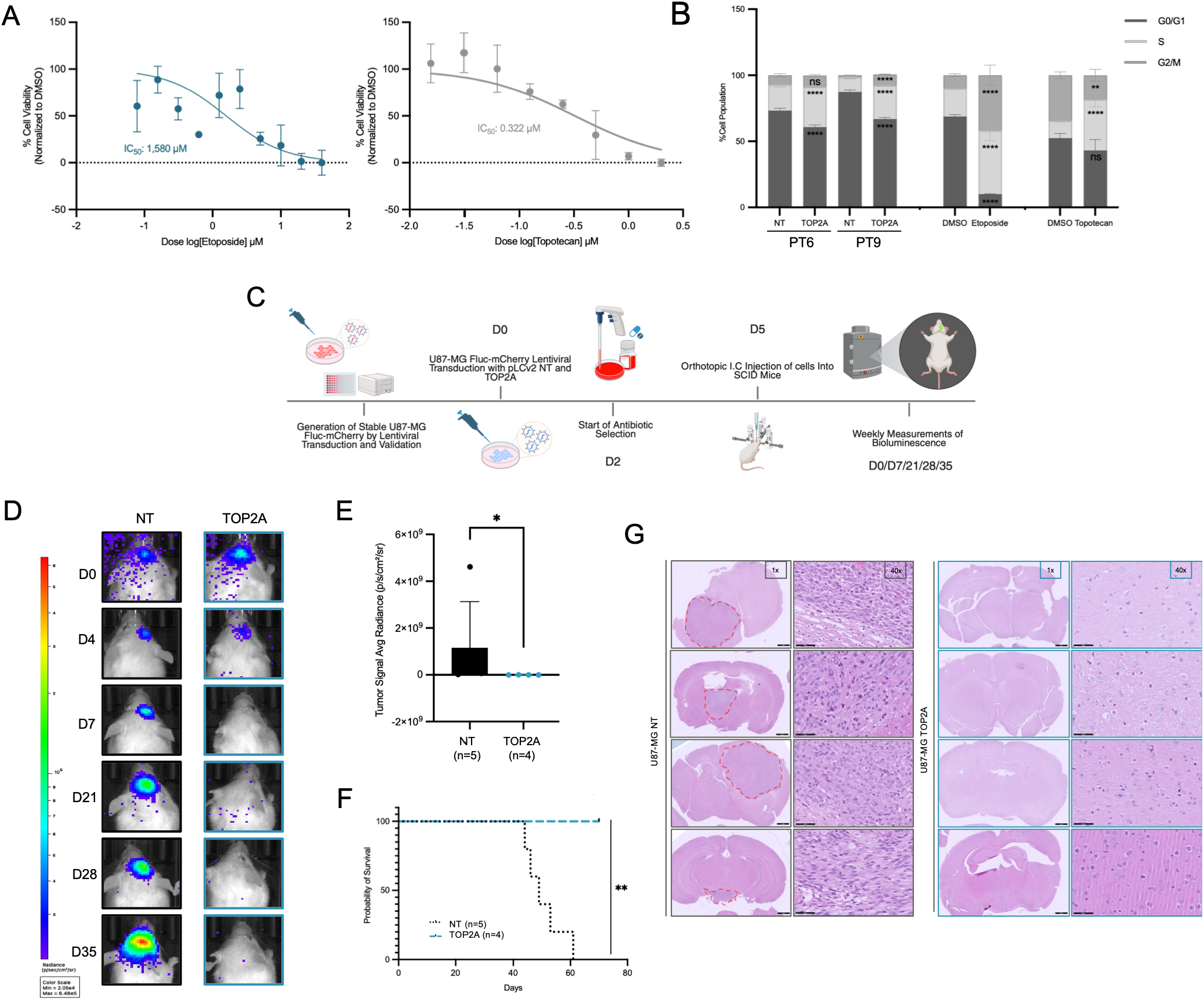
TOP2A depletion arrests cell cycle progression and abrogates orthotopic glioblastoma growth in vivo. **A)** Dose–response curves of U87-MG cells treated with etoposide and topotecan, demonstrating differential sensitivity under indicated experimental conditions. IC50 values were calculated based on normalized cell viability measurements. **B)** Cell cycle distribution analysis of U87-MG cells following TOP2A depletion and etoposide or topotecan treatment. Cells transduced with TOP2A-targeting sgRNAs or NT controls were treated with IC50 concentrations of etoposide or topotecan, and the proportions of cells in G0/G1, S, and G2/M phases were determined by flow cytometry. Stacked bar graphs represent the relative distribution of cell cycle phases under each experimental condition. Statistical analyses were performed using two-way ANOVA followed by Šídák’s multiple comparisons test to compare knockout and control groups, and drug treatment and DMSO groups within each cell cycle phase and time point. **C)** Schematic of orthotopic intracranial implantation experiment. U87-Fluc-mCherry cells stably expressing NT or TOP2A sgRNA were injected into SCID mice at day 5; bioluminescence measured weekly. **D)** Representative bioluminescence images at indicated timepoints. **E)** Quantification of tumor signal average radiance at day 35, the data was analyzed using an unpaired two-tailed Student’s *t*-test. **F)** Kaplan-Meier survival curves of SCID mice bearing orthotopic U87 Firefly-Luciferase-mCherry tumors transduced with non-targeting (NT, n=5) or TOP2A-targeting (n=4) sgRNA. **G)** Representative H&E-stained sections from NT and TOP2A KO tumors. Differences in overall survival between NT and TOP2A groups were analyzed using the log-rank (Mantel–Cox) test. Data are presented as mean ± SEM. Statistical significance was defined as ns, not significant; *p* < 0.05 (\**), p < 0.01 (**), p < 0.001 (\*\*\**), and *p < 0.0001 (****)*.

We next tested whether TOP2A is required for tumor growth in vivo. U87-MG cells stably expressing firefly luciferase and mCherry were transduced with NT or TOP2A sgRNA, antibiotic-selected, and implanted orthotopically into SCID mice, with tumor burden monitored weekly by bioluminescence imaging (**Fig. 6C**). Before implantation, TOP2A depletion in this reporter line was confirmed by qRT-PCR (**Supp. Fig. 7A**) and was accompanied by significantly reduced viability (**Supp. Fig. 7B**) and clonogenic capacity (**Supp. Fig. 7C**) in vitro, establishing an effective knockout at the time of injection. Tumor growth diverged markedly between groups. NT animals developed progressively expanding intracranial tumors, with bioluminescent signal throughout (**Fig. 6D**). Quantification confirmed significantly reduced tumor average radiance at day 35 (**Fig. 6E**). Critically, this translated into a survival benefit: all NT animals succumbed between approximately day 42 and day 62, while every TOP2A-knockout animal survived to the end of the study (**Fig. 6F**). Brain sections from NT animals contained large, well-demarcated, densely cellular tumor masses with the hypercellularity and nuclear atypia characteristic of glioblastoma, whereas TOP2A-knockout brains showed no comparable lesions and retained largely normal parenchymal architecture (**Fig. 6G**). Together, these results establish TOP2A as a central node linking chromosome segregation and DNA repair dependencies in glioblastoma, show that its genetic loss is phenocopied by clinically available topoisomerase inhibitors, and demonstrate that TOP2A depletion abrogates orthotopic tumor formation and significantly prolongs survival in vivo.

## DISCUSSION

Here we developed DDRKOL, a compact pathway-focused CRISPR/Cas9 knockout library targeting 819 DNA damage response genes and demonstrated its utility for systematically defining DNA repair dependencies in glioblastoma. By combining dense sgRNA representation with a manageable library size, DDRKOL provides an experimentally accessible alternative to genome-wide screening while maintaining sufficient resolution to prioritize functionally relevant genes within a biologically coherent pathway. Application of DDRKOL across two independent glioblastoma models identified a conserved set of shared vulnerabilities, with subsequent orthogonal validation establishing TOP2A, CDK1, XRCC6, and RAD21 as bona fide regulators of glioblastoma fitness. Collectively, these findings demonstrate that focused functional genomic approaches can identify clinically relevant genetic dependencies. The rationale underlying DDRKOL is distinct from that of genome-wide CRISPR libraries. Genome-scale collections are ideally suited for unbiased gene discovery, whereas pathway-focused libraries prioritize analytical depth within a defined biological process. Increasing guide representation from the four to six sgRNAs typically used in genome-wide libraries to ten independent guides per gene improves confidence in gene-level depletion estimates and facilitates discrimination between candidate genes with similar phenotypic effects [19,20,34]. At the same time, the substantially reduced library size lowers the cell numbers, sequencing depth, and experimental cost required for each screen, making biological replication, drug combination studies, and modifier screens considerably more practical. Although we applied DDRKOL to glioblastoma in the present study, its design is readily transferable to any Cas9-competent experimental system in which DNA damage response biology is of interest. We applied DDRKOL here to glioblastoma, but the library is not specific to this disease, and we return to its broader utility below.

Because the biological conclusions depend on library performance, the quality control data warrant explicit consideration. Sequencing of the plasmid pool recovered 8,969 of 8,970 designed sgRNAs, and representation was uniform, with Lorenz curves approximating the diagonal and unimodal count distributions across samples. The Gini coefficients obtained (0.261 in plasmid, 0.284 and 0.291 in cells) are above the ≤0.1 threshold proposed as an ideal for plasmid libraries [27], and we do not wish to overstate them; for a ten-guide library, however, the more informative question is whether individual genes were left under-represented, and the median relative minimum sgRNA count (0.48 in plasmid, 0.38 in both cell lines) exceeded the 0.3 threshold for acceptable per-gene coverage. Correlations against the plasmid pool (r = 0.758 and r = 0.750) indicate that complexity survived transduction and selection without bottlenecking. Internal controls provide independent support: Cas9 activity was confirmed by GFP reporter assay prior to screening, essential-gene sgRNAs depleted strongly while non-targeting and non-essential controls remained neutral [35], and pan-essential genes recovered here behaved consistently with their DepMap profiles [29]. Most substantively, all four prioritized candidates validated across orthogonal assays in two cell lines and in patient-derived models.

The rationale for interrogating the DDR follows from the mechanism of standard therapy. Radiotherapy and temozolomide both act by inducing DNA damage, and glioblastoma cells survive both by repairing that damage efficiently [3,6]. MGMT promoter methylation accounts for part of the variability in treatment response but not all of it [5], implying that additional repair factors contribute to the therapeutic ceiling. Consistent with this, dependencies concentrated in homologous recombination, nucleotide excision repair, and ATM/DSB signaling rather than distributing uniformly across the network. The two cell lines nonetheless differed substantially in hit number (106 in A172 versus 23 in U87-MG), a discrepancy that may reflect biological divergence or differences in editing efficiency, screen sensitivity, or proliferation rate; as we cannot presently distinguish these, we treated the twenty shared hits as the high-confidence set.

The individual knockout validations were not uniform in efficiency. Transcript levels in U87-MG were reduced to approximately 24–63% of control, whereas A172 was more variable, with TOP2A reduced to 15–20%, XRCC6 to only 56–66%, and one RAD21 guide producing no significant reduction; Western blotting reflected the same pattern. Two factors are relevant. First, validations were performed in polyclonal populations, such that each sample comprises a mixture of edited and unedited alleles, including in-frame indels that preserve function and can obscure genuine dependencies [36]. Second, and more consequentially, phenotypes were robust regardless: cells retaining appreciable residual protein still exhibited reduced viability, impaired colony formation, and progressive depletion in competition assays. Because complete null alleles are selected against during outgrowth of an essential gene knockout, surviving populations become enriched for hypomorphic alleles, and our measurements therefore represent a conservative estimate of the underlying dependency.

TOP2A emerged as the most compelling candidate, and the co-dependency analysis offers a mechanistic rationale. The genes most strongly correlated with TOP2A dependency in DepMap were components of the chromosome segregation machinery rather than DNA repair enzymes, including most of the condensin complex together with CENPA, ESPL1, KIF23, and RACGAP1 [29]. The concurrent appearance of TOP3A and RMI1 indicates a second axis concerned with resolving DNA topological intermediates. These observations position TOP2A at a decatenation node where chromosome segregation and topological stress resolution converge, a dependency difficult to buffer given that intact repair capacity cannot compensate for failure to separate replicated chromosomes. The cell cycle data support this interpretation, as both genetic depletion and pharmacological inhibition redistributed cells from G0/G1 into S and G2/M. Notably, TOP2A expression is highest in IDH-wildtype tumors, the most proliferative and least tractable glioma subtype.

The existence of clinically established topoisomerase-targeting agents represents both an opportunity and a constraint. Etoposide and topotecan reduced glioblastoma cell viability at sub- to low-micromolar concentrations here, yet neither has improved outcomes in this disease. The limiting factor appears to be delivery rather than target biology, as systemic administration is constrained by toxicity and limited blood-brain barrier penetration. This may be surmountable: chronic convection-enhanced delivery of topotecan into peritumoral brain was well tolerated in patients with recurrent glioblastoma and reduced proliferating tumor cell populations [37]. Isoform selectivity is a second consideration, as secondary malignancies associated with etoposide arise predominantly through TOP2β whereas cytotoxicity in transformed cells depends principally on TOP2α [38], supporting development of α-selective agents over existing pan-TOP2 inhibitors. Earlier phase 1b experience similarly demonstrated radiographic antitumor activity and prolonged survival with convection-enhanced topotecan at concentrations that were non-toxic to normal brain, providing clinical proof-of-concept that locoregional delivery can overcome an important pharmacological limitation of topoisomerase-directed therapy in malignant glioma [39].

The remaining validated candidates also illustrate the functional diversity of DNA damage response dependencies in glioblastoma. CDK1 behaved as anticipated, correlating with proliferation and adverse prognosis, with inhibitors already under evaluation in glioblastoma [13]. In contrast, RAD21 and XRCC6 revealed more subtle effects. RAD21 presented a more complex profile: knockout impaired fitness consistently across all models, yet elevated expression associated with prolonged survival in TCGA. Although RAD21 depletion consistently impaired cellular fitness, its elevated expression correlated with improved patient survival, emphasizing that transcriptional abundance and functional dependency capture distinct biological properties. XRCC6 showed the converse pattern, with no survival association or Ki-67 correlation and only a modest effect in the GBM8 viability assay despite reducing spheroid volume. Given its role in initiating non-homologous end joining [14], XRCC6 may contribute less to baseline proliferation than to survival following induced damage, positioning it as a radiosensitization candidate rather than a monotherapy target. Together, these observations underscore the importance of integrating functional genetic screening with expression-based analyses when prioritizing therapeutic targets. This interpretation is supported by the development of small-molecule Ku70/80 inhibitors that disrupt Ku-DNA binding and sensitize human glioblastoma cells to radiation at sub-cytotoxic concentrations [40], providing a direct pharmacological rationale for evaluating XRCC6 inhibition in combination with radiotherapy. RAD21 may likewise offer a combination strategy rather than an immediately druggable monotherapy target: although no direct RAD21 inhibitor is currently available, RAD21 silencing increased sensitivity to olaparib, rucaparib, and niraparib in ovarian cancer models by impairing double-strand break repair [41]. Because PARP inhibition has already demonstrated tumor penetration and clinical feasibility in recurrent glioblastoma, including in the phase I OPARATIC study of olaparib plus temozolomide [42], testing whether RAD21 loss or suppression creates a comparable PARP-inhibitor vulnerability in glioblastoma is a transitionally relevant next step.

Beyond the specific dependencies reported here, DDRKOL is a general-purpose resource, and its most valuable applications likely lie ahead. The DDR is dysregulated across essentially all solid and hematological malignancies, and the library can be deployed unchanged in any Cas9-competent model to identify tumor-type-specific repair vulnerabilities. Its more distinctive utility, however, is in modifier screening. Because the library is small enough to run in parallel arms at manageable cost, it can be screened under selective pressure to identify genes whose loss sensitizes cells to, or confers resistance against, a given therapy. Applied with temozolomide or irradiation, this would define the genes glioblastoma requires to survive treatment rather than the genes it requires to grow; these are unlikely to be identical sets, and the former is where clinical resistance is determined. The same design extends naturally to the expanding class of DDR-directed agents, including PARP, ATR, ATM, DNA-PK, CHK1, and WEE1 inhibitors, where rational combination partners and resistance mechanisms remain incompletely defined [43]. Screens of this kind could also nominate synthetic lethal interactions specific to repair deficiencies, an approach that has already proven clinically productive in homologous recombination-deficient tumors [43].

Several limitations should be acknowledged. The primary screens employed two established cell lines, one of which, U87-MG, differs genetically from its tumor of origin [44]; we addressed this by validating candidates in patient-derived spheroid models [32], though the caveat applies to the screen itself. The in vivo experiments used a single orthotopic model with limited group sizes and addressed whether TOP2A is required for tumor establishment rather than whether its inhibition is therapeutic in established disease. Finally, we have not yet applied DDRKOL under genotoxic selection, which, as outlined above, we consider the most immediate and informative extension of this work.

## CONCLUSIONS

This study establishes DDRKOL as a dedicated functional genomic resource for interrogating the DNA damage response in cancer, and applies it to define the DDR dependencies of glioblastoma. Parallel screens in two glioblastoma cell lines identified a shared, cell-line-independent core of dependencies concentrated in homologous recombination, nucleotide excision repair, and ATM/DSB signaling. From this set, TOP2A, CDK1, XRCC6, and RAD21 were validated across orthogonal assays and in patient-derived spheroid models. TOP2A proved the most therapeutically compelling: its depletion arrested cell cycle progression, and abolished orthotopic tumor formation while significantly prolonging survival in vivo.

The broader significance lies in the resource itself. The DDR is dysregulated across essentially all malignancies, and DDRKOL can be applied unchanged in any Cas9-competent model to identify tumor-type-specific vulnerabilities. Its most valuable application is likely modifier screening: applied under temozolomide, irradiation, or DDR-directed agents, it can define the genes tumors require to survive treatment rather than those required to grow, nominating rational combination partners and mechanisms of resistance. For a disease in which outcomes have not meaningfully improved in decades, mapping the repair pathways that permit survival of standard therapy is a necessary step toward better treatment.

## LIST OF ABBREVIATIONS

ATM: ataxia-telangiectasia mutated
ATR: ataxia telangiectasia and Rad3-related
CRISPR: clustered regularly interspaced short palindromic repeats
DDR: DNA damage response
DDRKOL: DNA Damage Response Knockout Library
DNA-PK: DNA-dependent protein kinase
DSB: double-strand break
EP: EndPoint
FITC: fluorescein isothiocyanate
FLuc: firefly luciferase
GBM: glioblastoma
GBMLGG: glioblastoma and lower-grade glioma
GeCKO: Genome-Scale CRISPR Knock-Out
GEPIA2: Gene Expression Profiling Interactive Analysis 2
GTEx: Genotype-Tissue Expression
IC50: half-maximal inhibitory concentration
IDH: isocitrate dehydrogenase
IP: InitialPoint
MAGeCK: Model-based Analysis of Genome-wide CRISPR/Cas9 Knockout
MGMT: O6-methylguanine-DNA methyltransferase
MOI: multiplicity of infection
MTT: 3-(4,5-dimethylthiazol-2-yl)-2,5-diphenyltetrazolium bromide
NHEJ: non-homologous end joining
NOD/SCID: non-obese diabetic/severe combined immunodeficiency
NT: non-targeting control
PARP: poly(ADP-ribose) polymerase
PI: propidium iodide
PT: post-transduction
qRT-PCR: quantitative reverse transcription polymerase chain reaction
RNAi: RNA interference
sgRNA: single-guide RNA
TCGA: The Cancer Genome Atlas
TOP1: DNA topoisomerase I
TOP2: DNA topoisomerase II

## DECLARATIONS

### Ethics approval and consent to participate

All animal experiments were conducted with the approval of the Koç University Animal Experiments Local Ethics Committee (HADYEK; approval no. 2020-007)

### Consent for publication

All authors have read and approved the final manuscript and consent to its publication.

### Availability of data and materials

The CRISPR screening NGS data generated in this study have been deposited in the NCBI Gene Expression Omnibus (GEO). Figures created with BioRender.com have been licensed for publication. (Fig. 1A: MG29ZV90IB, Fig. 2A: IR2A06GL1H, Fig. 4B: RS2A06GW5B, Fig. 4G: ZW2A06TI2F, Fig. 6C: DX2A014TOQ).

### Authors’ contributions

Study design: I.S.E., A.K., A.C.A., T.B.O.; Methodology: I.S.E., A.K., A.C.A., T.B.O.; Data generation and

analysis: C.B.O., I.K., A.O., A.H.D.K., I.K.; Data interpretation: C.B.O, I.K., A.O., T.B.O.; Drafted the manuscript: I.K., C.B.O., T.B.O.; Approved final manuscript: all authors.

## Acknowledgements

The authors gratefully acknowledge the use of the services and facilities of the Koç University Research Center for Translational Medicine (KUTTAM). We also thank Dr. Ozlem Yedier-Bayram for her valuable scientific input.

