## Supplementary Material for "DDRKOL: A Focused CRISPR Library for Systematic Identification of DNA Damage Response Dependencies in Glioblastoma"

### SUPPLEMENTARY FIGURES

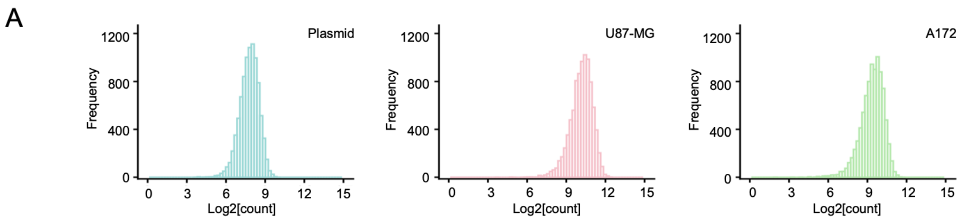


**Supplementary Figure 1. sgRNA counts for plasmid and cell lines.** Frequency histograms of $log_2$-transformed normalized sgRNA counts for the plasmid library (teal), U87-MG (pink), and A172 (green) cell lines, demonstrating successful lentiviral transduction and uniform library representation across samples.

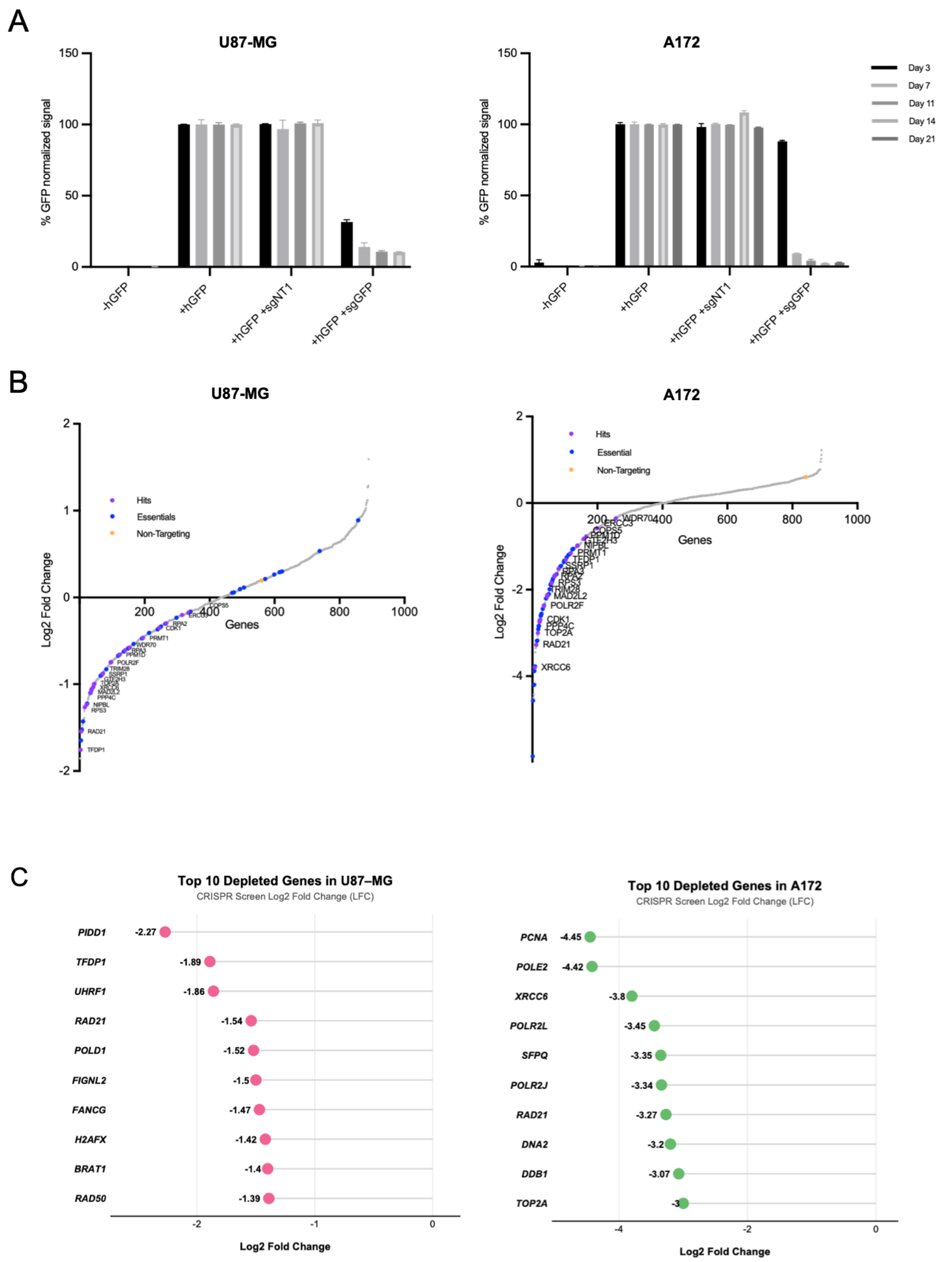


**Supplementary Figure 2. Cas9 efficiency and screen readouts in glioblastoma cell lines. A)** GFP-based Cas9-activity assay results for U87-MG and A172 cell lines. **B)** Waterfall plots illustrating the distribution of gene-level CRISPR scores ($log_2$ fold change, LFC) for U87-MG (left) and A172 (right) cell lines. Genes are ranked by their LFC between the InitialPoint and EndPoint (PDL 15). Individual data points represent screen hits (purple), pan-essential positive controls (blue), non-targeting controls (orange). **C)** Lollipop plots displaying the top 10 most significantly depleted genes in U87-MG (pink) and A172 (green) screens, ranked by their $log_2$ fold change. Values to the right of each dot indicate the specific LFC for each candidate.

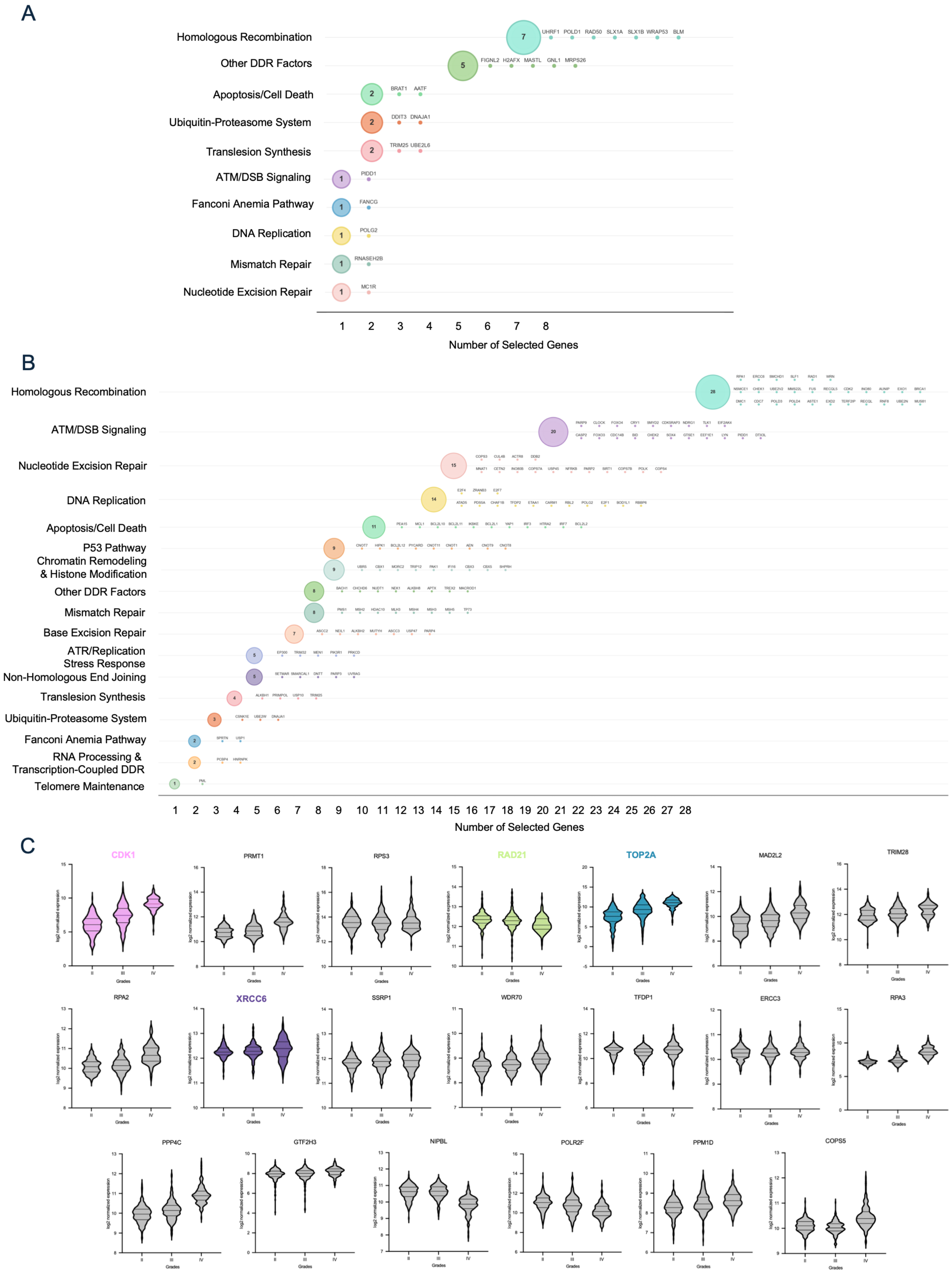
A
Homologous Recombination
Other DDR Factors
Apoptosis/Cell Death
Ubiquitin-Proteasome System
Translesion Synthesis
ATM/DSB Signaling
Fanconi Anemia Pathway
DNA Replication
Mismatch Repair
Nucleotide Excision Repair
UHRF1 POLD1 RAD50 SLX1A SLX1B WRAP53 BLM
FIGNL2 H2AFX MASTL GNL1 MRPS26
BRAT1 AATF
DDIT3 DNAJA1
TRIM25 UBE2L6
PIDD1
FANCG
POLG2
RNASEH2B
MC1R
1 2 3 4 5 6 7 8
Number of Selected Genes
B
Homologous Recombination
ATM/DSB Signaling
Nucleotide Excision Repair
DNA Replication
Apoptosis/Cell Death
P53 Pathway
Chromatin Remodeling & Histone Modification
Other DDR Factors
Mismatch Repair
Base Excision Repair
ATR/Replication Stress Response
Non-Homologous End Joining
Translesion Synthesis
Ubiquitin-Proteasome System
Fanconi Anemia Pathway
RNA Processing & Transcription-Coupled DDR
Telomere Maintenance
1 2 3 4 5 6 7 8 9 10 11 12 13 14 15 16 17 18 19 20 21 22 23 24 25 26 27 28
Number of Selected Genes
C
CDK1
PRMT1
RPS3
RAD21
TOP2A
MAD2L2
TRIM28
RPA2
XRCC6
SSRP1
WDR70
TFDP1
ERCC3
RPA3
PPP4C
GTF2H3
NIPBL
POLR2F
PPM1D
COPS5
log2 normalized expression
Grades
II III IV

**Supplementary Figure 3. Pathway analysis and clinical relevance of CRIPSR screen hits. A-B)** Pathway-level distribution of screen hits identified in U87-MG **(A)** and A172 **(B)** models. Bubbles indicate the number of selected genes per functional category, with individual gene members listed to the right. **C)** Violin plots showing the mRNA expression levels ($\log_2$ normalized) of the 21 common screen hits across histological glioma grades (Grade II, III, and IV) from the TCGA dataset.

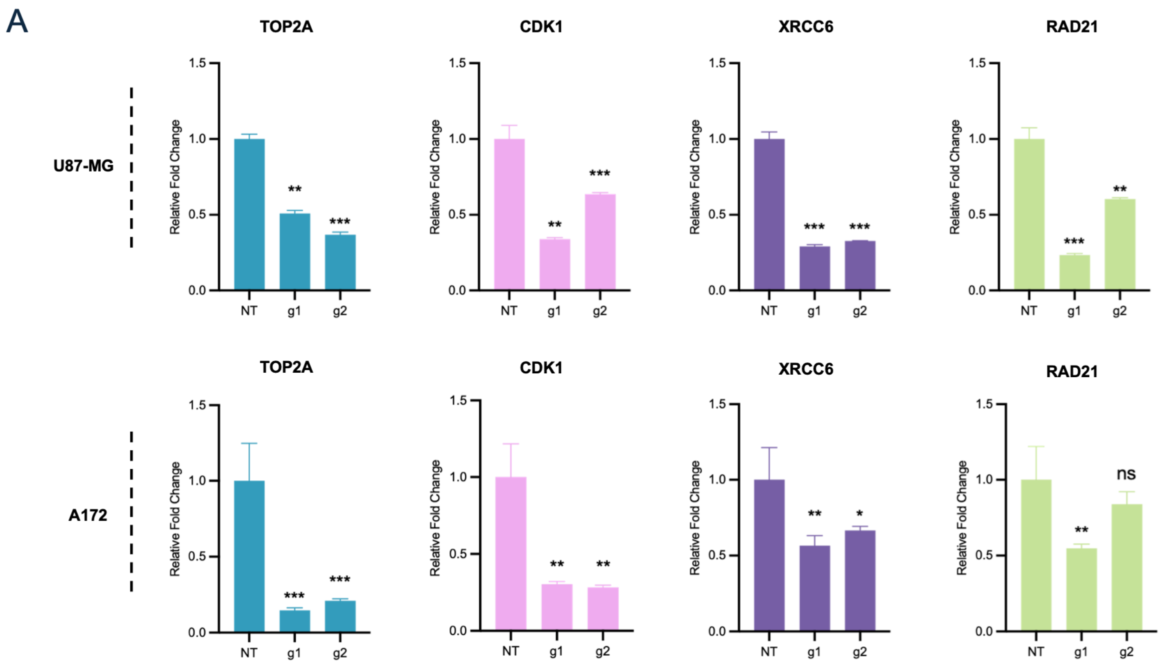


**Supplementary Figure 4. Quantitative RT-PCR (qRT-PCR) validation of mRNA depletion** following CRISPR-mediated knockout using two independent sgRNAs (g1 and g2) in U87-MG and A172 cells. Relative fold change is normalized to non-targeting (NT) controls. Statistical significance was determined by one-way ANOVA (*p < 0.05, **p < 0.01, ***p < 0.001; ns: non-significant).

A

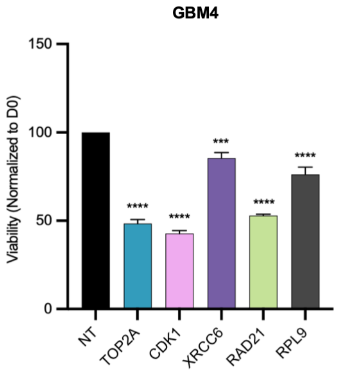


**Supplementary Figure 5. Cell viability in primary GBM4 spheroids.** Cell Titer Glo (CTG) assay for PT9 in GBM4 patient-derived cells following indicated knockouts. Statistical comparisons to NT; *p<0.05, **p<0.01, ***p<0.001, ****p<0.0001.

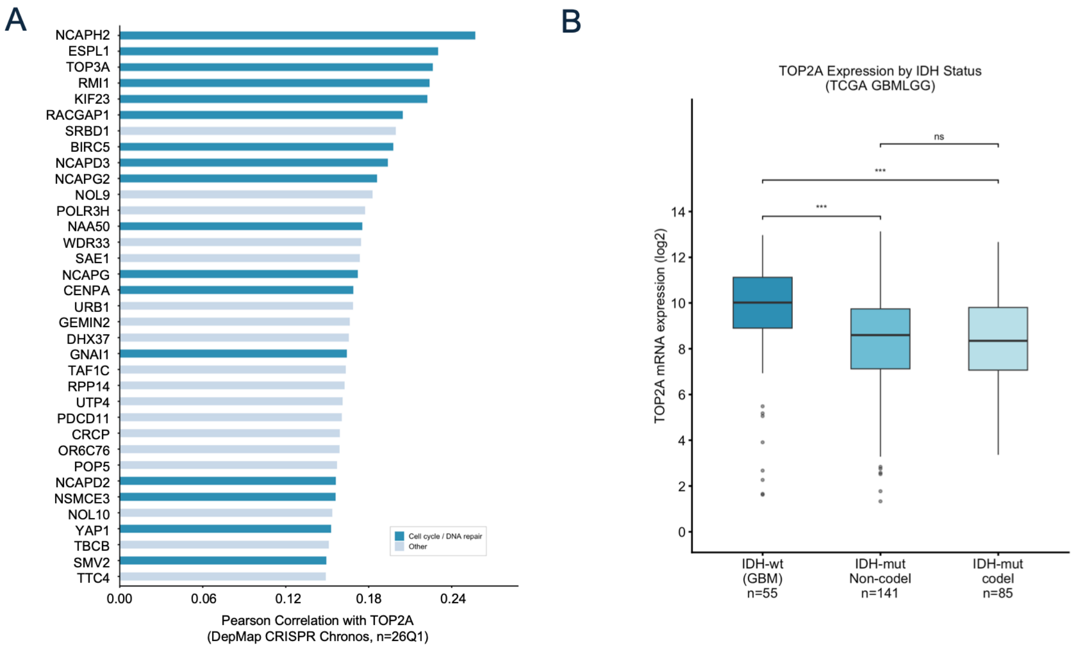


**Supplementary Figure 6. Analysis of co-dependency and clinical relevance of TOP2A. A)** DepMap CRISPR co-dependency analysis of TOP2A across cancer cell lines. Pearson correlation coefficients between TOP2A and all other genes were obtained from the DepMap Public 26Q1 CRISPR (Chronos) dataset. The top 35 positively correlated genes were ranked based on co-dependency scores and functionally annotated using Gene Ontology (GO) terms associated with cell cycle and DNA repair pathways. Genes linked to GO:0007049, GO:0000278, GO:0006281, or GO:0006974 were classified as "Cell cycle / DNA repair," while remaining genes were categorized as "Other." **B)** TOP2A mRNA expression by IDH status (TCGA GBMLGG): IDH-wt GBM (n=55), IDH-mut non-codeletion (n=141), IDH-mut codeletion (n=85).

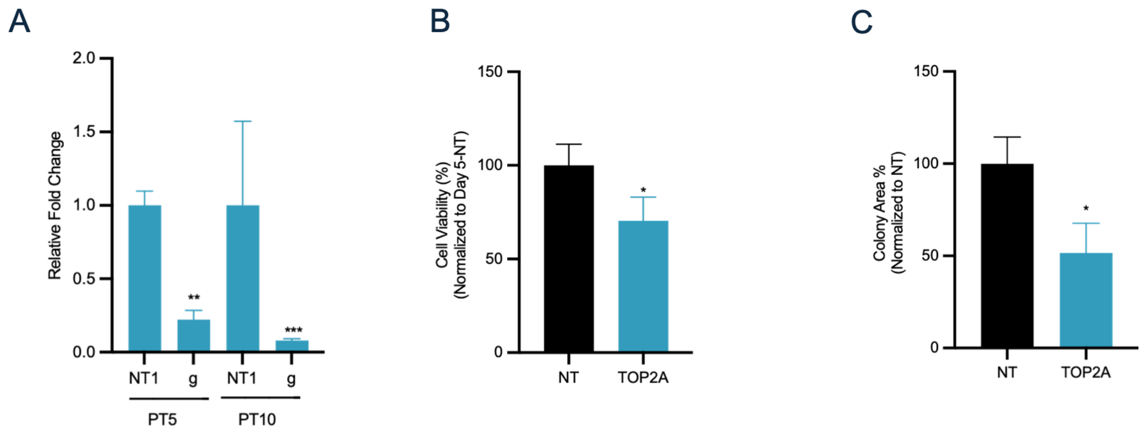


**Supplementary Figure 7. Validation of *in vivo* compatible U87-Fluc-mCherry cell line. A)** Quantitative RT-PCR analysis confirming significant depletion of *TOP2A* mRNA levels in U87-Fluc-mCherry (FMC) cells at PT5 and PT10. Relative fold change is normalized to the non-targeting control (NT). **B)** MTT viability assay in U87-FMC cells at day 5 post-selection, demonstrating a significant reduction in cellular proliferation following *TOP2A* knockout compared to the NT control. **C)** Quantitative analysis of colony formation assays in U87-FMC cells. Bars represent the percentage of colony area normalized to NT controls, indicating that *TOP2A* depletion profoundly impairs the long-term clonogenic capacity of the cell population intended for orthotopic xenografting. *p<0.05. Statistical significance was determined by Student's t-test (*p < 0.05, **p < 0.01, ***p < 0.001). All error bars represent SD from three independent replicates.

#### SUPPLEMENTARY INFORMATION

##### Cas9 Activity Assay

To determine the Cas9 activity of U87MG and A172, Flow Cytometry was used. GBM cells were transduced with pLentiCas9-blast and Hygro-GFP lentiviruses in this order with low MOI (0.4). Non-transduced cells were selected with Blasticidin (10 µg/ml) and Hygromycin (200 µg/ml) respectively. U87MG/A172-Cas9-GFP cells were later transduced with either non-targeting control vector (NT) or GFP targeting vector (T1) and selected with puromycin (2 µg/ml). %GFP was measured with flow cytometry every 3 days.

##### Co-dependency Analysis

TOP2A co-dependency analysis used pre-computed Pearson correlation scores from the DepMap Public 26Q1 CRISPR (Chronos) dataset, retrieved via the DepMap gene co-dependency portal. The top 35 positively correlated genes were selected for visualization. Functional annotation was performed using the gprofiler2 R package (v2.x): genes were queried against GO:BP, KEGG, and Reactome database, and genes intersecting any of a curated set of cell cycle or DNA repair–related term IDs (including GO:0007049, GO:0000278, GO:0006281, GO:0006974, and related terms) were classified as "Cell cycle / DNA repair"; remaining genes were classified as "Other." Bar plots were generated using ggplot2 at 300 DPI.

#### SUPPLEMENTARY TABLES

##### Supplementary Table 2:

| Primer | Sequence (5' to 3') |
|---|---|
| PLG_Ext_Forward | AATGGACTATCATATGCTTACCGTAACTTGAAAGTATTTCG |
| PLG_Ext_Reverse | CTTTAGTTTGTATGTCTGTTGCTATTATGTCTACTATTCTTTCC |

##### Supplementary Table 3:

| Primer | Sequence (5' to 3') | Stagger |
|---|---|---|
| Forward_stag1 | AATGATACGGCGACCACCGAGATCTACACTCTTTCCCTACACGACGCTCTTCCGATCTAtcttgtggaaaggacgaaacaccg | A |
| Forward_stag2 | AATGATACGGCGACCACCGAGATCTACACTCTTTCCCTACACGACGCTCTTCCGATCTCTtcttgtggaaaggacgaaacaccg | CT |
| Forward_stag3 | AATGATACGGCGACCACCGAGATCTACACTCTTTCCCTACACGACGCTCTTCCGATCTGACtcttgtggaaaggacgaaacaccg | GAC |
| Forward_stag4 | AATGATACGGCGACCACCGAGATCTACACTCTTTCCCTACACGACGCTCTTCCGATCTTCGAtcttgtggaaaggacgaaacaccg | TCGA |
| Forward_stag5 | AATGATACGGCGACCACCGAGATCTACACTCTTTCCCTACACGACGCTCTTCCGATCTATCGCtcttgtggaaaggacgaaacaccg | ATCGC |
| Forward_stag6 | AATGATACGGCGACCACCGAGATCTACACTCTTTCCCTACACGACGCTCTTCCGATCTGTCAGAtcttgtggaaaggacgaaacaccg | GTCAGA |
| Forward_stag7 | AATGATACGGCGACCACCGAGATCTACACTCTTTCCCTACACGACGCTCTTCCGATCTTGCATCGtcttgtggaaaggacgaaacaccg | TGCATCG |
| Forward_stag8 | AATGATACGGCGACCACCGAGATCTACACTCTTTCCCTACACGACGCTCTTCCGATCTCTACGTGAtcttgtggaaaggacgaaacaccg | CTACGTGA |
| Forward_stag9 | AATGATACGGCGACCACCGAGATCTACACTCTTTCCCTACACGACGCTCTTCCGATCTACTCGTGATtcttgtggaaaggacgaaacaccg | ACTCGTGAT |

##### Supplementary Table 4:

| Primer | Sequence (5' to 3') | Index |
|---|---|---|
| rev_index2 | CAAGCAGAAGACGGCATACGAGATACATCGGTGACTGGAGTTCAGACGTGTGCTCTTCCGATCTtctactattctttcccctgcactgt | ACATCG |
| rev_index3 | CAAGCAGAAGACGGCATACGAGATGCCTAAGTGACTGGAGTTCAGACGTGTGCTCTTCCGATCTtctactattctttcccctgcactgt | GCCTAA |
| rev_index4 | CAAGCAGAAGACGGCATACGAGATTGGTCAGTGACTGGAGTTCAGACGTGTGCTCTTCCGATCTtctactattctttcccctgcactgt | TGGTCA |
| rev_index5 | CAAGCAGAAGACGGCATACGAGATCACTGTGTGACTGGAGTTCAGACGTGTGCTCTTCCGATCTtctactattctttcccctgcactgt | CACTGT |
| rev_index6 | CAAGCAGAAGACGGCATACGAGATATTGGCGTGACTGGAGTTCAGACGTGTGCTCTTCCGATCTtctactattctttcccctgcactgt | ATTGGC |
| rev_index7 | CAAGCAGAAGACGGCATACGAGATGATCTGGTGACTGGAGTTCAGACGTGTGCTCTTCCGATCTtctactattctttcccctgcactgt | GATCTG |
| rev_index8 | CAAGCAGAAGACGGCATACGAGATTCAAGTGTGACTGGAGTTCAGACGTGTGCTCTTCCGATCTtctactattctttcccctgcactgt | TCAAGT |
| rev_index9 | CAAGCAGAAGACGGCATACGAGATCTGATCGTGACTGGAGTTCAGACGTGTGCTCTTCCGATCTtctactattctttcccctgcactgt | CTGATC |
| rev_index10 | CAAGCAGAAGACGGCATACGAGATAAGCTAGTGACTGGAGTTCAGACGTGTGCTCTTCCGATCTtctactattctttcccctgcactgt | AAGCTA |
| rev_index11 | CAAGCAGAAGACGGCATACGAGATGTAGCCGTGACTGGAGTTCAGACGTGTGCTCTTCCGATCTtctactattctttcccctgcactgt | GTAGCC |

| Primer | Sequence (5' to 3') | Index |
|---|---|---|
| rev_index12 | CAAGCAGAAGACGGCATACGAGATTACAAGGTGACTGGAGTTCAGACGTGTGCTCTTCCGATCTtctactattctttcccctgcactgt | TACAAG |
| rev_index13 | CAAGCAGAAGACGGCATACGAGATTTGACTGTGACTGGAGTTCAGACGTGTGCTCTTCCGATCTtctactattctttcccctgcactgt | TTGACT |
| rev_index14 | CAAGCAGAAGACGGCATACGAGATGGAACTGTGACTGGAGTTCAGACGTGTGCTCTTCCGATCTtctactattctttcccctgcactgt | GGAACT |
| rev_index15 | CAAGCAGAAGACGGCATACGAGATTGACATGTGACTGGAGTTCAGACGTGTGCTCTTCCGATCTtctactattctttcccctgcactgt | TGACAT |
| rev_index16 | CAAGCAGAAGACGGCATACGAGATGGACGGGTGACTGGAGTTCAGACGTGTGCTCTTCCGATCTtctactattctttcccctgcactgt | GGACGG |
| rev_index17 | CAAGCAGAAGACGGCATACGAGATCTCTACGTGACTGGAGTTCAGACGTGTGCTCTTCCGATCTtctactattctttcccctgcactgt | CTCTAC |
| rev_index18 | CAAGCAGAAGACGGCATACGAGATGCGGACGTGACTGGAGTTCAGACGTGTGCTCTTCCGATCTtctactattctttcccctgcactgt | GCGGAC |
| rev_index19 | CAAGCAGAAGACGGCATACGAGATTTTCACGTGACTGGAGTTCAGACGTGTGCTCTTCCGATCTtctactattctttcccctgcactgt | TTTCAC |
| rev_index20 | CAAGCAGAAGACGGCATACGAGATGGCCACGTGACTGGAGTTCAGACGTGTGCTCTTCCGATCTtctactattctttcccctgcactgt | GGCCAC |
| rev_index21 | CAAGCAGAAGACGGCATACGAGATCGAAACGTGACTGGAGTTCAGACGTGTGCTCTTCCGATCTtctactattctttcccctgcactgt | CGAAAC |
| rev_index22 | CAAGCAGAAGACGGCATACGAGATCGTACGGTGACTGGAGTTCAGACGTGTGCTCTTCCGATCTtctactattctttcccctgcactgt | CGTACG |
| rev_index23 | CAAGCAGAAGACGGCATACGAGATCCACTCGTGACTGGAGTTCAGACGTGTGCTCTTCCGATCTtctactattctttcccctgcactgt | CCACTC |
| rev_index24 | CAAGCAGAAGACGGCATACGAGATGCTACCGTGACTGGAGTTCAGACGTGTGCTCTTCCGATCTtctactattctttcccctgcactgt | GCTACC |
| rev_index25 | CAAGCAGAAGACGGCATACGAGATATCAGTGTGACTGGAGTTCAGACGTGTGCTCTTCCGATCTtctactattctttcccctgcactgt | ATCAGT |
| rev_index26 | CAAGCAGAAGACGGCATACGAGATGCTCATGTGACTGGAGTTCAGACGTGTGCTCTTCCGATCTtctactattctttcccctgcactgt | GCTCAT |
| rev_index27 | CAAGCAGAAGACGGCATACGAGATAGGAATGTGACTGGAGTTCAGACGTGTGCTCTTCCGATCTtctactattctttcccctgcactgt | AGGAAT |
| rev_index28 | CAAGCAGAAGACGGCATACGAGATCTTTTGGTGACTGGAGTTCAGACGTGTGCTCTTCCGATCTtctactattctttcccctgcactgt | CTTTTG |
| rev_index29 | CAAGCAGAAGACGGCATACGAGATTAGTTGGTGACTGGAGTTCAGACGTGTGCTCTTCCGATCTtctactattctttcccctgcactgt | TAGTTG |
| rev_index30 | CAAGCAGAAGACGGCATACGAGATCCGGTGGTGACTGGAGTTCAGACGTGTGCTCTTCCGATCTtctactattctttcccctgcactgt | CCGGTG |
| rev_index31 | CAAGCAGAAGACGGCATACGAGATATCGTGGTGACTGGAGTTCAGACGTGTGCTCTTCCGATCTtctactattctttcccctgcactgt | ATCGTG |
| rev_index32 | CAAGCAGAAGACGGCATACGAGATTGAGTGGTGACTGGAGTTCAGACGTGTGCTCTTCCGATCTtctactattctttcccctgcactgt | TGAGTG |
| rev_index33 | CAAGCAGAAGACGGCATACGAGATCGCCTGGTGACTGGAGTTCAGACGTGTGCTCTTCCGATCTtctactattctttcccctgcactgt | CGCCTG |
| rev_index34 | CAAGCAGAAGACGGCATACGAGATGCCATGGTGACTGGAGTTCAGACGTGTGCTCTTCCGATCTtctactattctttcccctgcactgt | GCCATG |
| rev_index35 | CAAGCAGAAGACGGCATACGAGATAAAATGGTGACTGGAGTTCAGACGTGTGCTCTTCCGATCTtctactattctttcccctgcactgt | AAAATG |
| rev_index36 | CAAGCAGAAGACGGCATACGAGATTGTTGGGTGACTGGAGTTCAGACGTGTGCTCTTCCGATCTtctactattctttcccctgcactgt | TGTTGG |
| rev_index37 | CAAGCAGAAGACGGCATACGAGATATTCCGGTGACTGGAGTTCAGACGTGTGCTCTTCCGATCTtctactattctttcccctgcactgt | ATTCCG |
| rev_index38 | CAAGCAGAAGACGGCATACGAGATAGCTAGGTGACTGGAGTTCAGACGTGTGCTCTTCCGATCTtctactattctttcccctgcactgt | AGCTAG |
| rev_index39 | CAAGCAGAAGACGGCATACGAGATGTATAGGTGACTGGAGTTCAGACGTGTGCTCTTCCGATCTtctactattctttcccctgcactgt | GTATAG |
| rev_index40 | CAAGCAGAAGACGGCATACGAGATTCTGAGGTGACTGGAGTTCAGACGTGTGCTCTTCCGATCTtctactattctttcccctgcactgt | TCTGAG |

| Primer | Sequence (5' to 3') | Index |
|---|---|---|
| rev_index41 | CAAGCAGAAGACGGCATACGAGATGTCGTCGTGACTGGAGTTCAGACGTGTGCTCTTCCGATCTtctactattctttcccctgcactgt | GTCGTC |
| rev_index42 | CAAGCAGAAGACGGCATACGAGATCGATTAGTGACTGGAGTTCAGACGTGTGCTCTTCCGATCTtctactattctttcccctgcactgt | CGATTA |
| rev_index43 | CAAGCAGAAGACGGCATACGAGATGCTGTAGTGACTGGAGTTCAGACGTGTGCTCTTCCGATCTtctactattctttcccctgcactgt | GCTGTA |
| rev_index44 | CAAGCAGAAGACGGCATACGAGATATTATAGTGACTGGAGTTCAGACGTGTGCTCTTCCGATCTtctactattctttcccctgcactgt | ATTATA |
| rev_index45 | CAAGCAGAAGACGGCATACGAGATGAATGAGTGACTGGAGTTCAGACGTGTGCTCTTCCGATCTtctactattctttcccctgcactgt | GAATGA |
| rev_index46 | CAAGCAGAAGACGGCATACGAGATTCGGGAGTGACTGGAGTTCAGACGTGTGCTCTTCCGATCTtctactattctttcccctgcactgt | TCGGGA |
| rev_index47 | CAAGCAGAAGACGGCATACGAGATCTTCGAGTGACTGGAGTTCAGACGTGTGCTCTTCCGATCTtctactattctttcccctgcactgt | CTTCGA |
| rev_index48 | CAAGCAGAAGACGGCATACGAGATTGCCGAGTGACTGGAGTTCAGACGTGTGCTCTTCCGATCTtctactattctttcccctgcactgt | TGCCGA |

#### Supplementary Table 5:

| Target Gene | Sequence (5' to 3') | DDRKOL ID |
|---|---|---|
| CDK1_1_F | CACCGGATCTCCAGAAGTATTGCTG | DDRKoL_1183 |
| CDK1_1_R | AAACCAGCAATACTTCTGGAGATCC | |
| CDK1_2_F | CACCGTATACCAAAATAGAGAAAAT | DDRKoL_1186 |
| CDK1_2_R | AAACATTTTCTCTATTTTGGTATAC | |
| RAD21_1_F | CACCGGATCGTGAGATAATGAGAGA | DDRKoL_5956 |
| RAD21_1_R | AAACTCTCTCATTATCTCACGATCC | |
| RAD21_2_F | CACCGGTGTAATTTAGAGAGCAGCG | DDRKoL_5953 |
| RAD21_2_R | AAACCGCTGCTCTCTAAATTACACC | |
| RPL9_1_F | CACCGAGGGCTTCCGTTACAAGATG | DDRKoL_6622 |
| RPL9_1_R | AAACCATCTTGTAACGGAAGCCCTC | |
| RPL9_2_F | CACCGATGACTACAAATAGTCCGAA | DDRKoL_6623 |
| RPL9_2_R | AAACTTCGGACTATTTGTAGTCATC | |
| TOP2A_1_F | CACCGAGCATTGTAAAGATGTATCG | DDRKoL_7871 |
| TOP2A_1_R | AAACCGATACATCTTTACAATGCTC | |
| TOP2A_2_F | CACCGTAATTCCACAGAACCAATGT | DDRKoL_7876 |
| TOP2A_2_R | AAACACATTGGTTCTGTGGAATTAC | |
| XRCC6_1_F | CACCGGGTGATCTCCGAGATACAGG | DDRKoL_8702 |
| XRCC6_1_R | AAACCCTGTATCTCGGAGATCACCC | |
| XRCC6_2_F | CACCGAGGTTCGCGCCAAGGAGACC | DDRKoL_8706 |
| XRCC6_2_R | AAACGGTCTCCTTGGCGCGAACCTC | |
| NT_F | ACGGAGGCTAAGCGTCGCAA | |
| NT_R | TTGCGACGCTTAGCCTCCGT | |
| T1_F | CACCGTGAACCGCATCGAGCTGAA | |
| T1_R | AAACTTCAGCTCGATGCGGTTCAC | |

#### Supplementary Table 6:

| Name | Company/Catalog Number | Species | Molecular Weight | Web Page |
|---|---|---|---|---|
| anti-Alpha Tubulin | Elabscience/E-AB-20069 | Rabbit | 50 kDa | https://www.elabscience.com/viewpdf-26175-Elabscience-E-AB-20069.pdf |
| anti-CDK1 | PTG 19532-1-AP-20UL | Rabbit | 30-34 kDa | https://www.ptglab.com/products/CDC2-Specific-Antibody-19532-1-AP.htm |
| Anti-Histone H3 (D1H2) XP (R) | Cell Signaling/4499S | Rabbit | 17 kDa | https://www.cellsignal.com/products/primary-antibodies/histone-h3-d1h2-xp-rabbit-mab/4499?srsltid=AfmBOooKxz-7wpvVVNGv4TQGdK3kXi8GPWSmQU3JTJCoT52Q1ybmVsNI |
| anti-RAD21 | PTG 27071-1-AP-20UL | Rabbit | 120 kDa | https://www.ptglab.com/products/RAD21-Antibody-27071-1-AP.htm |
| anti-TOP2A | PTG 24641-1-AP-20UL | Rabbit | 174 kDa | https://www.ptglab.com/products/TOP2A-Antibody-24641-1-AP.htm |
| anti-XRCC6 (Ku70) | PTG 66607-1-IG-20UL | Mouse | 70 kDa | https://www.ptglab.com/products/KU70%2CXRCC6-Antibody-66607-1-Ig.htm |
| Goat anti-Mouse IgG H&L (HRP) | Abcam/ab97023 | - | - | https://www.abcam.com/en-us/products/secondary-antibodies/goat-mouse-igg-h-l-hrp-ab97023 |
| Goat anti-Rabbit IgG H&L (HRP) | Abcam/ab6721 | - | - | https://www.abcam.com/en-us/products/secondary-antibodies/goat-rabbit-igg-h-l-hrp-ab6721 |

#### Supplementary Table 7:

| Target Gene | Sequence (5' to 3') |
|---|---|
| CDK1_qPCR_F | GGAAACCAGGAAGCCTAGCATC |
| CDK1_qPCR_R | GGATGATTCAGTGCCATTTTGCC |
| RAD21_qPCR_F | GTGGAAAGAGACAGGAGGAGTAG |
| RAD21_qPCR_R | AGGTCTTCTGGTACAAGCGGTG |
| RPL9_qPCR_F | GCACAGTTATCGTGAAGGGC |
| RPL9_qPCR_R | TTACCCCACCATTTGTCAACC |
| TOP2A_qPCR_F | GTGGCAAGGATTCTGCTAGTCC |
| TOP2A_qPCR_R | ACCATTCAGGCTCAACACGCTG |
| XRCC6_qPCR_F | TCATGGCAACTCCAGAGCAG |
| XRCC6_qPCR_R | AACCTTGGGCAATGTCAGGT |
